# Proprioceptive limit detectors mediate sensorimotor control of the *Drosophila* leg

**DOI:** 10.1101/2025.05.15.654260

**Authors:** Brandon G. Pratt, Chris J. Dallmann, Grant M. Chou, Igor Siwanowicz, Sarah Walling-Bell, Andrew Cook, Anne Sustar, Anthony Azevedo, John C. Tuthill

## Abstract

Many animals possess mechanosensory neurons that fire when a limb nears the limit of its physical range, but the function of these proprioceptive limit detectors remains poorly understood. Here, we investigate a class of proprioceptors on the *Drosophila* leg called hair plates. Using calcium imaging in behaving flies, we find that a hair plate on the fly coxa (CxHP8) detects the limits of anterior leg movement. Reconstructing CxHP8 axons in the connectome, we found that they are wired to excite posterior leg movement and inhibit anterior leg movement. Consistent with this connectivity, optogenetic activation of CxHP8 neurons elicited posterior postural reflexes, while silencing altered the swing-to-stance transition during walking. Finally, we use comprehensive reconstruction of peripheral morphology and downstream connectivity to predict the function of other hair plates distributed across the fly leg. Our results suggest that each hair plate is specialized to control specific sensorimotor reflexes that are matched to the joint limit it detects. They also illustrate the feasibility of predicting sensorimotor reflexes from a connectome with identified proprioceptive inputs and motor outputs.

## Introduction

Animals rely on proprioception to sense and coordinate movements of the body. One key function of the proprioceptive system is to detect when a body part has reached the limits of its normal range. In vertebrates, including mammals, the extremes of joint position are detected by low-threshold Ruffini endings and Pacinian corpuscles embedded within joint capsules (Burgess and Clark, 1969). In arthropods, including insects, joint limits may be detected by hair plates, fields of small, stiff mechanosensory hairs positioned at cuticular folds within joints (Pringle, 1938). Compared to proprioceptors that encode limb position and movement, such as vertebrate muscle spindles and invertebrate chordotonal organs, less is known about the physiology and behavioral function of proprioceptors that detect joint limits (Tuthill and Azim, 2018).

Early recordings from joint receptors in cats found that single afferents fired in a phasic or tonic manner near the limit of the range of joint movement (Burgess and Clark, 1969). For much of the 20^th^ century, it was believed that these receptors were the primary source of our conscious sense of body position, or kinesthesia (Proske and Gandevia, 2012). However, humans retain their kinesthetic sense after joint capsules are removed following hip replacement surgery (Grigg et al., 1973). This and other evidence (Clark et al., 1979) suggests that muscle spindles are the primary proprioceptors underlying kinesthesia in mammals. Although joint receptors may not be required for kinesthesia, they could still play a role in unconscious, reflexive control of movement, a hypothesis which has received less attention since the focus shifted to muscle spindles (Proske and Gandevia, 2012).

In insects and other invertebrates, feedback from proprioceptive limit detectors has been shown to initiate stabilizing reflexes that sculpt motor patterns and protect the limbs from injury (Bässler, 1977). For example, classic work in the cockroach showed that hair plate neurons at a proximal leg joint monosynaptically excite extensor motor neurons and indirectly inhibit flexor motor neurons, restricting the range of joint flexion (Pearson et al., 1976; Wong and Pearson, 1976). The ablation of this hair plate caused the corresponding leg to overstep and collide with the more anterior leg during walking, indicating that limit detection is important for controlling phase transitions of the step cycle (Wong and Pearson, 1976). Past work has identified specific sensorimotor reflexes mediated by insect hair plates, but it has been difficult to synthesize into a more comprehensive understanding of the underlying feedback circuits and their function. Like many proprioceptors, hair plates are distributed throughout the animal’s body, which raises the possibility that each sensory organ is specialized for sensing and controlling specific movements.

With emerging connectomic wiring maps (Azevedo et al., 2024; Dorkenwald et al., 2024; Takemura et al., 2024; Winding et al., 2022) and genetic tools to target specific cell types (Jenett et al., 2012), the fruit fly, *Drosophila*, has recently emerged as an important model system to investigate neural mechanisms of limb proprioception. We and others have used connectomics (Lee et al., 2024; Marin et al., 2024), electrophysiology (Agrawal et al., 2020), calcium imaging (Chen et al., 2021; Dallmann et al., 2023; Mamiya et al., 2023, 2018), and behavior (Akitake et al., 2015; Cheng et al., 2010; Chockley et al., 2022; Isakov et al., 2016; Mendes et al., 2013; Pratt et al., 2024), to investigate the function of proprioceptors in the fly’s femoral chordotonal organ (FeCO). The FeCO contains subtypes of mechanosensory neurons that detect the position, movement, and vibration of the fly femur-tibia joint, in a manner analogous to vertebrate muscle spindles (Mamiya et al., 2023, 2018). However, compared to the FeCO, little is currently known about the anatomy, physiology, and behavioral function of hair plates on the *Drosophila* leg.

The six *Drosophila* legs have 214 hair plate mechanosensory neurons that are clustered into 42 hair plates (**Figure S1A**) (Hodgkin and Bryant, 1979; Kuan et al., 2020; Schubiger, 1968). Investigating the physiology and function of each hair plate, one-by-one, would be experimentally intractable, even in the tiny fruit fly. Instead, we start by studying one specific hair plate on the coxa of the fly’s front legs (CxHP8) using *in vivo* calcium imaging (**Figure 1**), connectomics (**Figure 2**), and genetic manipulations during behavior (**Figures 3, 4**). We find that the sensory tuning and motor reflex action of CxHP8 are well predicted by its external morphology and downstream connectivity to leg motor neurons. We therefore extend these sensorimotor reflex predictions to other hair plates positioned at different locations on the leg (**Figure 5**). Overall, our results provide insight into the behavioral function of proprioceptive limit detectors and illustrate a new approach for comprehensive analysis of distributed sensorimotor systems that can be scaled to other animals.

## Results

### CxHP8 neurons detect the limits of coxa rotation and adduction

On the fly’s front leg, three hair plates are located at the junction between the coxa (Cx) and thorax (**Figure 1A**). These hair plates (CxHP8, CxHP3, CxHP4) wrap the joint along the anterior-posterior axis. We screened Split-GAL4 driver lines to identify lines that label coxa hair plates (**Table S1**). The most specific and complete line we found (Oh et al., 2021) labeled CxHP8 neurons on the front and middle legs (6 and 3 out of 8 cells, respectively). Using this specific driver line, which we refer to as CxHP8-GAL4 (**Figure 1B**; **Figure S1B**), we first investigated the joint angle encoding properties of CxHP8 neurons.

**Figure 1.**
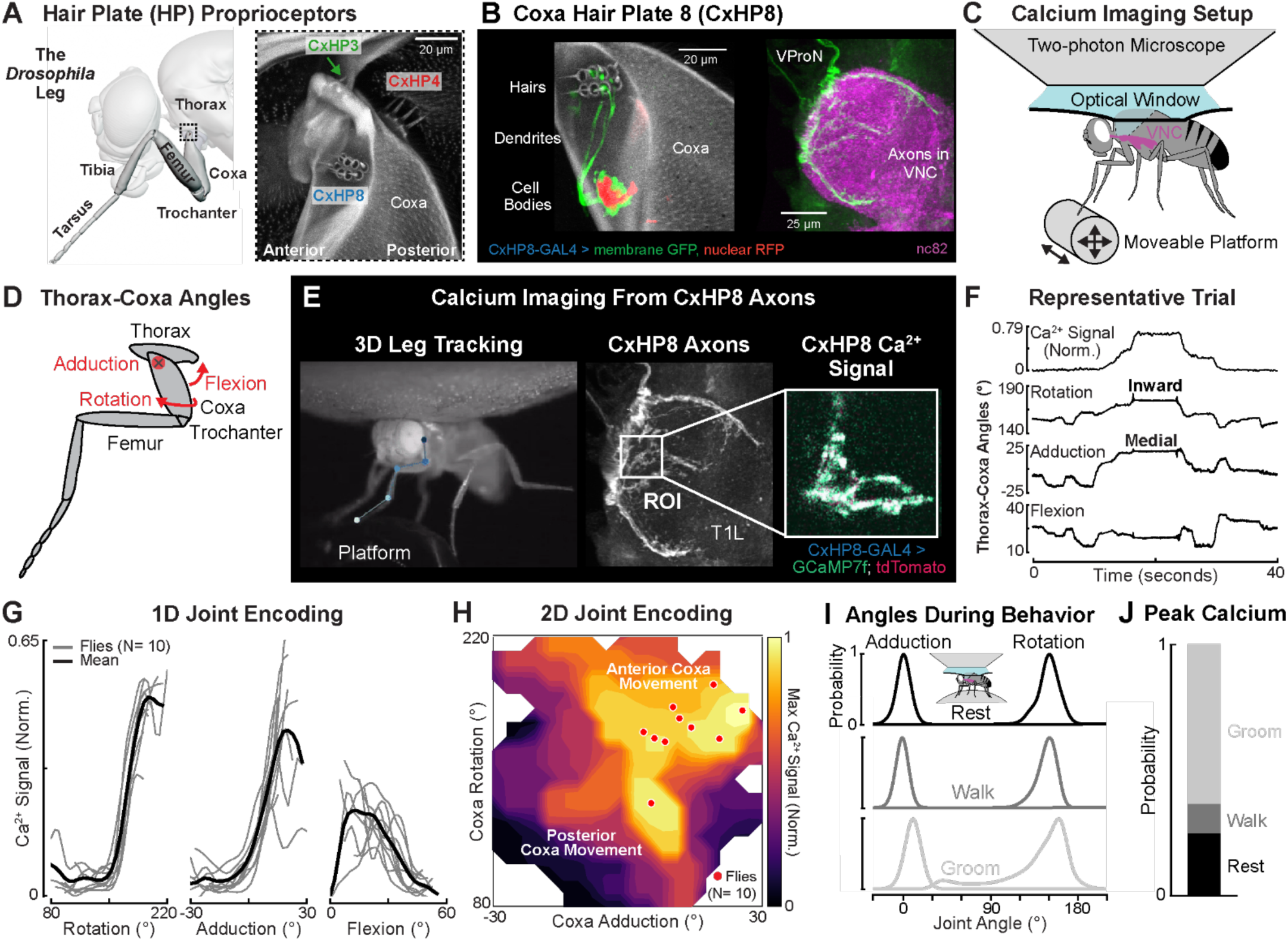
CxHP8 neurons encode the anterior limits of thorax-coxa joint angles. (**A**) Left: schematic rendering of the fly’s front leg. Right: a confocal image of the three hair plates, CxHP3, CxHP4, and CxHP8, at the thorax-coxa joint (black dotted box) of the front leg. (**B**) Confocal images of genetically labeled CxHP8 neurons (6 of 8) in the coxa of the front leg and ventral nerve cord (VNC). CxHP8 (autofluorescence: gray) consists of 8 cuticular hairs located on the coxa. Each hair is innervated by a single mechanosensory neuron (GFP: green), which projects through the ventral prothoracic nerve (VProN) into the VNC (nc82: magenta). The cell bodies of CxHP8 neurons are labeled in red (redstinger). (**C**) Schematic of the setup used to measure calcium signals from CxHP8 axons as the left front leg was passively moved on a platform. (**D**) Schematic of the rotation, adduction, and flexion thorax-coxa joint angles. (**E**) Example recording of calcium activity of CxHP8 axons. Joint angles were quantified by tracking the joints of the left front leg in 3D. In addition to GCaMP7f (green), CxHP8 axons expressed tdTomato (red) for motion correction and signal normalization. (**F**) Representative trial showing the normalized calcium signal for CxHP8 axons, the thorax-coxa rotation, adduction, and flexion angles while the left front leg was passively moved. Calcium activity was normalized between 0 and 1, where 1 represents the greatest calcium activity in the dataset. (**G**) Normalized calcium activity of CxHP8 axons with respect to thorax-coxa rotation, adduction, and flexion angles. Black line: bin averaged mean calcium activity across all flies; Gray lines: bin averaged mean calcium activity for each fly. (**H**) Contour plot showing the maximum 2D calcium activity across all flies with respect to thorax-coxa rotation and adduction. Red dots: the rotation and adduction angles at which the normalized calcium activity was maximum for each fly. (**I**) Adduction and rotation joint angle normalized probability densities across flies during rest (black), walking (gray), and front leg grooming (light gray). The inset shows a schematic of the calcium imaging setup used to determine CxHP8 activity during behavior. (**J**) Probability of flies grooming, walking, or resting during peak calcium activity. See also **Figure S1 and Videos 1-3**.

We used a previously described setup (Dallmann et al., 2023) that combines 2-photon calcium imaging with high-speed video to simultaneously record the activity of CxHP8 axons and 3D leg joint kinematics in CxHP8-GAL4 flies expressing GCaMP7f and tdTomato (**Figure 1C**). To overcome the slow temporal dynamics of GCaMP7f relative to the dynamics of active leg movement in flies, we passively (and slowly) moved the left front leg using a manually controlled 3-axis platform (**Video 1**). We then used DeepLabCut (Mathis et al., 2018) and Anipose (Karashchuk et al., 2021) to compute the 3 angles corresponding to the rotation, adduction, and flexion of the front leg thorax-coxa joint (**Figure 1D**). Increases in rotation, adduction, and flexion angles correspond to the coxa rotating inward, translating medially, and flexing, respectively (**Figure S1C**). Our goal in these experiments was to determine the relationship between thorax-coxa joint angles and CxHP8 calcium activity (**Figure 1E**).

Based on the anterior position of CxHP8 at the thorax-coxa joint, we hypothesized that its hairs would be deflected during anterior and medial movements, and inward rotations of the coxa (**Video 2**). A representative trial indeed shows a peak calcium signal when the coxa was inwardly rotated and adducted (**Figure 1F**). Across flies, we found similar tuning to coxa rotation and adduction (**Figure 1G**). Although with more variability, CxHP8 axons showed higher activity when the coxa was extended. Because coxa rotation and adduction produced strong calcium responses, we next investigated how the two angles jointly influenced the activity of CxHP8 neurons. We found that the combination of both adduction and inward rotation resulted in maximal calcium activity of CxHP8 axons (**Figure 1H**). Altogether, our results show that CxHP8 neurons detect the limits of anterior leg movement.

### CxHP8 neurons are active across behaviors

We next sought to determine which behaviors were accompanied by the extreme anterior coxa movements that drive CxHP8 activity. We used a similar imaging setup (Dallmann et al., 2023) but replaced the platform with an air-supported ball and allowed the fly to spontaneously behave (**Video 3**). We found that resting, walking, and grooming produced rotation and adduction coxa joint angles within the range that CxHP8 neurons were active (**Figure 1I**). As illustrated by a representative trace, calcium activity increased during grooming and walking compared to adjacent resting states (**Figure S1D**). Peak calcium activity was largest for front leg grooming bouts, likely because of the sustained deflection of CxHP8 cuticular hairs. Thresholding peak calcium activity (**Figure 1E**), we found CxHP8 neurons were active across behaviors (**Figure 1J**), particularly during extreme adduction and rotation angles (**Figure S1F**).

### Connectomics predict that CxHP8 neurons drive posterior leg movement

Using connectomics, we next reconstructed and analyzed hair plate sensorimotor circuits to predict the role of limit detection in leg motor control. Some hair plate axons were previously reconstructed in an electron microscopy (EM) dataset of a male adult nerve cord (MANC) (Marin et al., 2024). However, specific hair plate axons could not be identified in MANC due to poor reconstruction quality. We therefore reconstructed hair plate axons within an EM dataset of the Female Adult Nerve Cord (FANC)(Azevedo et al., 2024; Phelps et al., 2021). We first used previous annotations (Phelps et al., 2021) to identify and proofread all CxHP8 axons projecting into the left leg neuromere (**Figure 2A**). We identified the peripheral location of each hair plate axon in the FANC connectome using a combination of genetic driver lines, axonal morphology, axonal projection patterns, and an x-ray holographic nano-tomography dataset of the fly leg (Kuan et al., 2020) (See Methods for details). For example, CxHP8 cells are identifiable in the X-ray dataset because they are the only hair plate axons that project into the VNC through the ventral prothoracic nerve (VProN).

**Figure 2.**
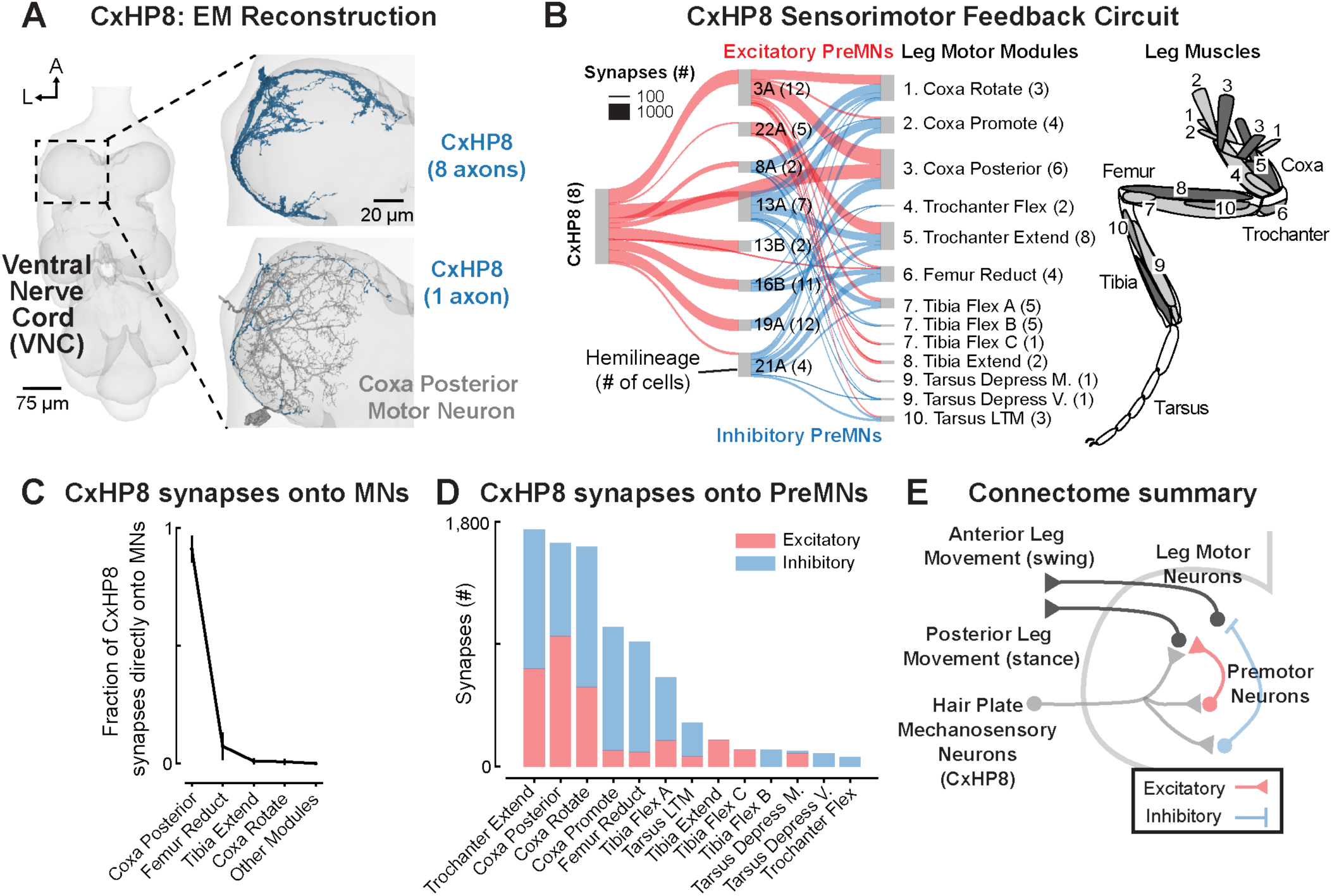
Synaptic connectivity suggests that CxHP8 neurons connect to a reflex circuit that drives posterior leg movement. (**A**) We reconstructed all CxHP8 axons within the left front leg neuromere of a *Drosophila* female VNC (FANC) electron microscopy volume. (**B**) CxHP8 axons synapse onto premotor neurons, identified by their hemilineage, as well as motor neurons grouped into motor modules (Azevedo et al., 2024). Red and blue lines signify excitatory and inhibitory connections, respectively. The strength of the connection (width of the line) is the number of synapses all neurons within a cell class make onto a downstream cell class. Only premotor and motor neurons that received an average of 4 or greater synapses per neuron from an upstream cell class were analyzed (i.e., CxHP8 input threshold: 8 cells * 4 synapses). The number of cells in each cell class included in this analysis is indicated within parentheses. The numbers in the leg schematic on the right illustrate the muscles belonging to leg motor modules 1-10. (**C**) CxHP8 neurons primarily synapse onto motor neurons within the coxa posterior motor module. Error bars are standard deviation. (**D**) Relative number of excitatory and inhibitory synapses between premotor neurons that receive input from CxHP8 neurons and motor neurons within motor modules. Red and blue bars indicate excitatory and inhibitory connections, respectively. (**E**) Summary schematic showing the circuit through which CxHP8 neurons are predicted to drive posterior leg movement. See also **Figure S2**.

The entire FANC dataset has been automatically segmented into neurons using convolutional neural networks (Azevedo et al., 2024), but many cells still required manual proofreading. We proofread and annotated cells downstream of CxHP8 neurons and found that CxHP8 axons allocate 75% of their output synapses (2,279 ± 482 synapses per CxHP8 axon) to premotor (interneurons that synapse on motor neurons: 58%) and motor (17%) neurons. This percentage is significantly higher than that of exteroceptive leg sensory neurons, but similar to FeCO proprioceptors (Lee et al., 2024), suggesting that hair plates are primarily involved in leg motor control.

The 69 motor neurons controlling the fly’s left front leg are organized into 14 motor modules – motor neurons within a module receive common presynaptic input and control the same joint (Azevedo et al., 2024). Premotor neurons in FANC have also been annotated based on their developmental hemilineage (Lesser et al., 2024), from which it is possible to infer the neurotransmitter released by each cell (acetylcholine, GABA, or glutamate). We first examined the connectivity from CxHP8 axons to premotor neurons to leg motor neurons (**Figure 2B**). CxHP8 neurons provide strong monosynaptic input to motor neurons within the coxa posterior module (**Figure 2B,C**). They also connect indirectly, via excitatory and inhibitory premotor neurons, to multiple other motor modules across the leg (**Figure 2C**, **Figure S2A**). CxHP8’s strongest inhibitory connections are onto the coxa promote module, an antagonist of the coxa posterior module (**Figure 2B**). We also found extensive recurrent connectivity within and across premotor neurons, as well as between CxHP8 axons and 19A premotor neurons (**Figure S2B**). Overall, CxHP8 neurons are positioned to provide feedback to multiple motor modules, but with the strongest excitation to motor neurons driving posterior leg movement and inhibition to motor neurons driving anterior movements. These connectivity patterns lead us to hypothesize that activity in CxHP8 neurons drives posterior leg movement when the coxa reaches its anterior limit. For example, CxHP8 feedback could contribute to the swing-to-stance transition during walking (**Figure 2E**).

### CxHP8 activation drives posterior leg movement and silencing alters swing-to-stance transitions during walking

We next tested whether CxHP8 activation contributes to posterior leg movements, as predicted by their synaptic connectivity to leg motor neurons. As in prior studies (Agrawal et al., 2020; Azevedo et al., 2020), we used a setup that enables spatiotemporally precise optogenetic activation or silencing of neurons in tethered flies behaving on an air-supported ball (**Figure 3A**). Using six high-speed cameras, we reconstructed full 3D leg joint kinematics using DeepLabCut (Mathis et al., 2018) and Anipose (Karashchuk et al., 2021). To optogenetically activate CxHP8 neurons, we genetically expressed ChrimsonR within the CxHP8 neurons and illuminated the thorax-coxa joint of the left front leg with a red laser. In a representative standing fly, we observed that its left front leg moved posteriorly shortly after the onset of the laser, which was accompanied by outward rotation, lateral movement, and flexion of the coxa (**Figure 3B,C**). We observed the same changes in thorax-coxa joint angles across standing flies (**Figure 3D**). These joint angle changes were accompanied by posterior, lateral, and downward displacement of the distal tarsus (**Figure 3E**). Overall, the activation of CxHP8 neurons produced posterior and lateral movement of the stimulated leg, without impacting the other legs.

**Figure 3.**
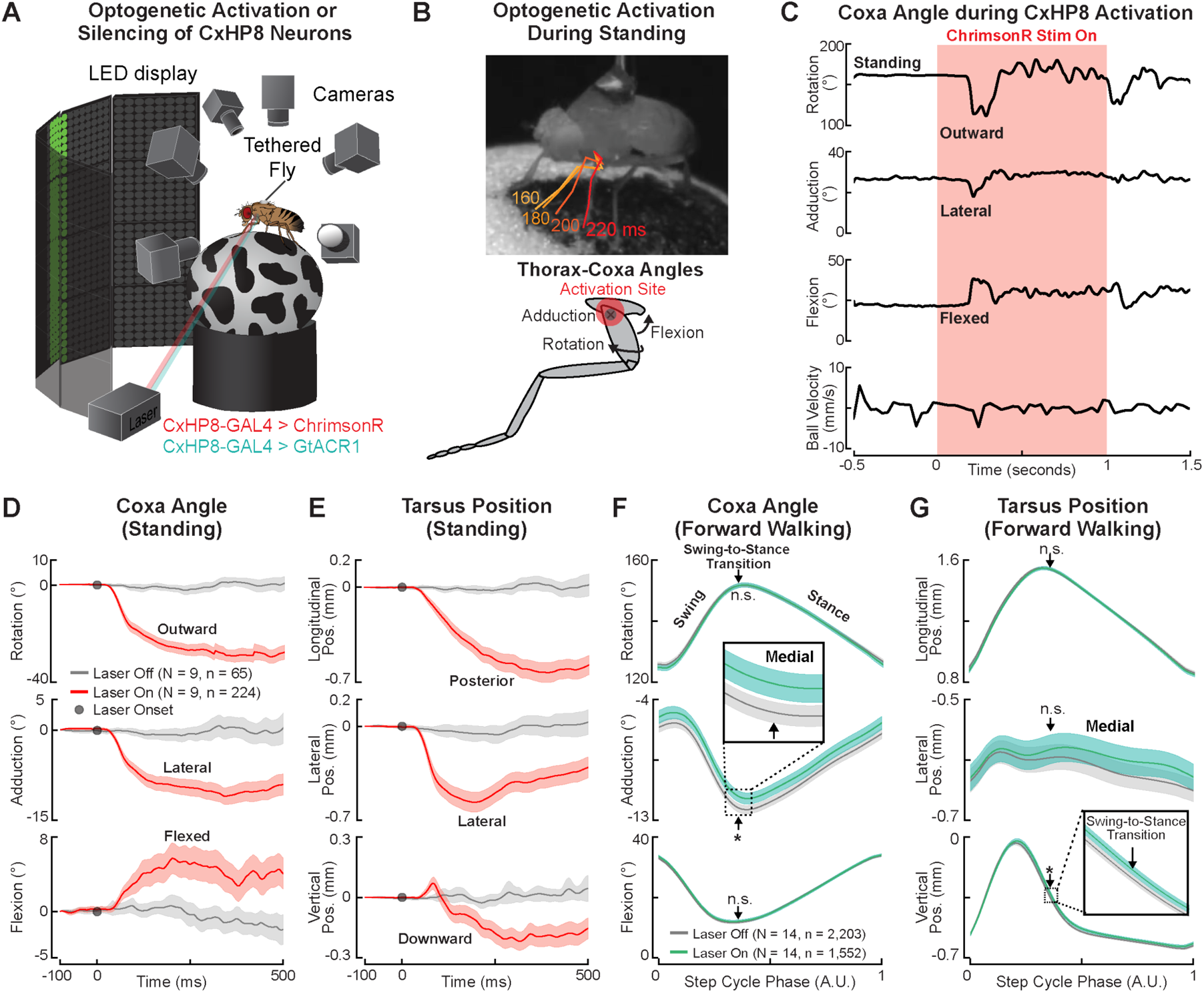
CxHP8 neurons drive posterior leg movement to mediate the swing-to-stance transition during walking. (**A**) Schematic of the optogenetic activation and silencing setup. 6 high-speed cameras (300 fps) recorded the behavior of a tethered fly on a ball and a laser focused on the left front leg was used to excite or inhibit CxHP8 neurons expressing the light-gated ion channel ChrimsonR (red) or GtACR1 (green). Trials were 2 s in duration and the laser was presented for 1 s during experimental trials and absent during control trials. (**B**) Top: example trial in which CxHP8 activation resulted in outward rotation, lateral movement, and flexion of the coxa in a standing fly. The amount of time, in ms, passed after the onset of the laser is displayed in the same color as the corresponding skeletonized leg. Bottom: the rotation, adduction, and flexion thorax-coxa joint angles. (**C**) For the same trial shown in **B**, the thorax-coxa rotation, adduction, and flexion angles, and the ball velocity in response to the optogenetic activation of CxHP8 neurons. (**D**) Optogenetic activation (red) of CxHP8 neurons in standing flies resulted in significant outward rotation, lateral displacement, and flexion of the coxa compared to trials in which the laser remained off (gray). Shaded region: 95% confidence interval; N: number of flies; n: number of trials. (**E**) Optogenetic activation (red) of CxHP8 neurons in standing flies also resulted in significant lateral and posterior movements of the tarsus compared to trials in which the laser remained off. The tarsus also moved downward along the ball during optogenetic activation. Shaded region: 95% confidence interval; N: number of flies; n: number of trials. (**F**) Optogenetic silencing (green) of CxHP8 neurons in flies walking forward produced a significant (*: p < 0.025) medial movement of the coxa near the swing-to-stance transition compared to forward walking steps when the laser was off (gray). Arrow: the average swing-to-stance transition of the step cycle; Shaded region: 95% confidence interval; N: number of flies; n: number of steps. A t-test with a Bonferroni correction of 2 was used to statistically compare the distributions of joint angles at the swing-to-stance transition. (**G**) Optogenetic silencing (green) of CxHP8 neurons in flies walking forward also resulted in a more medial and significantly higher (*: p < 0.025) tarsus position following the swing-to-stance transition compared to forward walking steps when the laser was off (gray). A t-test with a Bonferroni correction of 2 was used to statistically compare the distributions of tarsus positions at the swing-to-stance transition. See also **Figure S3**.

We next optogenetically silenced CxHP8 neurons through the expression of a light-gated chloride channel (GtACR1) during spontaneous behaviors. When CxHP8 feedback was disrupted during forward walking, the thorax-coxa adduction angle of the left front leg was slightly but significantly more medial, particularly at the swing-to-stance transition of the step cycle (**Figure 3F**). We did not observe medial coxa displacements in control flies that experienced the same laser stimulation (**Figure S3A**). Optogenetic silencing of CxHP8 neurons also subtly but significantly altered the tarsus position at the swing-to-stance transition (**Figure 3G**). Thus, the lack of CxHP8 activity appeared to cause the coxa to slightly overshoot the normal kinematic range associated with the swing-to-stance transition, leading to a medial displacement of the entire leg. We also assessed the role of CxHP8 feedback in left (**Figure S3B**) and right turns (**Figure S3C**), and front leg grooming (**Figure S3D**). Although joint angles were not affected during left turns, we found that the coxa was significantly more extended and rotated inward at the swing-to-stance transition when CxHP8 neurons were silenced during right turns. The coxa was also significantly more extended and adducted in flies with CxHP8 neurons silenced during front leg grooming.

### CxHP8 facilitates the swing-to-stance transition and controls resting posture in untethered flies

Although convenient for delivering targeted optogenetic stimuli and tracking joint kinematics, flies walking in the tethered configuration have slightly different walking kinematics than untethered flies (Pratt et al., 2024). To test whether CxHP8 feedback is important for the control of swing-to-stance transitions in untethered walking flies, we used a linear treadmill system (Pratt et al., 2024). We genetically expressed an inward rectifying potassium channel, Kir 2.1, to chronically hyperpolarize CxHP8 neurons. As flies walked on the treadmill, driven at a belt speed of 10 mm/s, we recorded their movement at 200 fps with 5 high-speed cameras (**Figure 4A**). From the high-speed videos, we reconstructed the 3D positions of the head, thorax, abdomen, and each leg tip using DeepLabCut (Mathis et al., 2018) and Anipose (Karashchuk et al., 2021). We then used these reconstructed 3D positions to quantify kinematics and posture during walking and rest periods, respectively (see Methods for details; **Figure 4B**).

**Figure 4.**
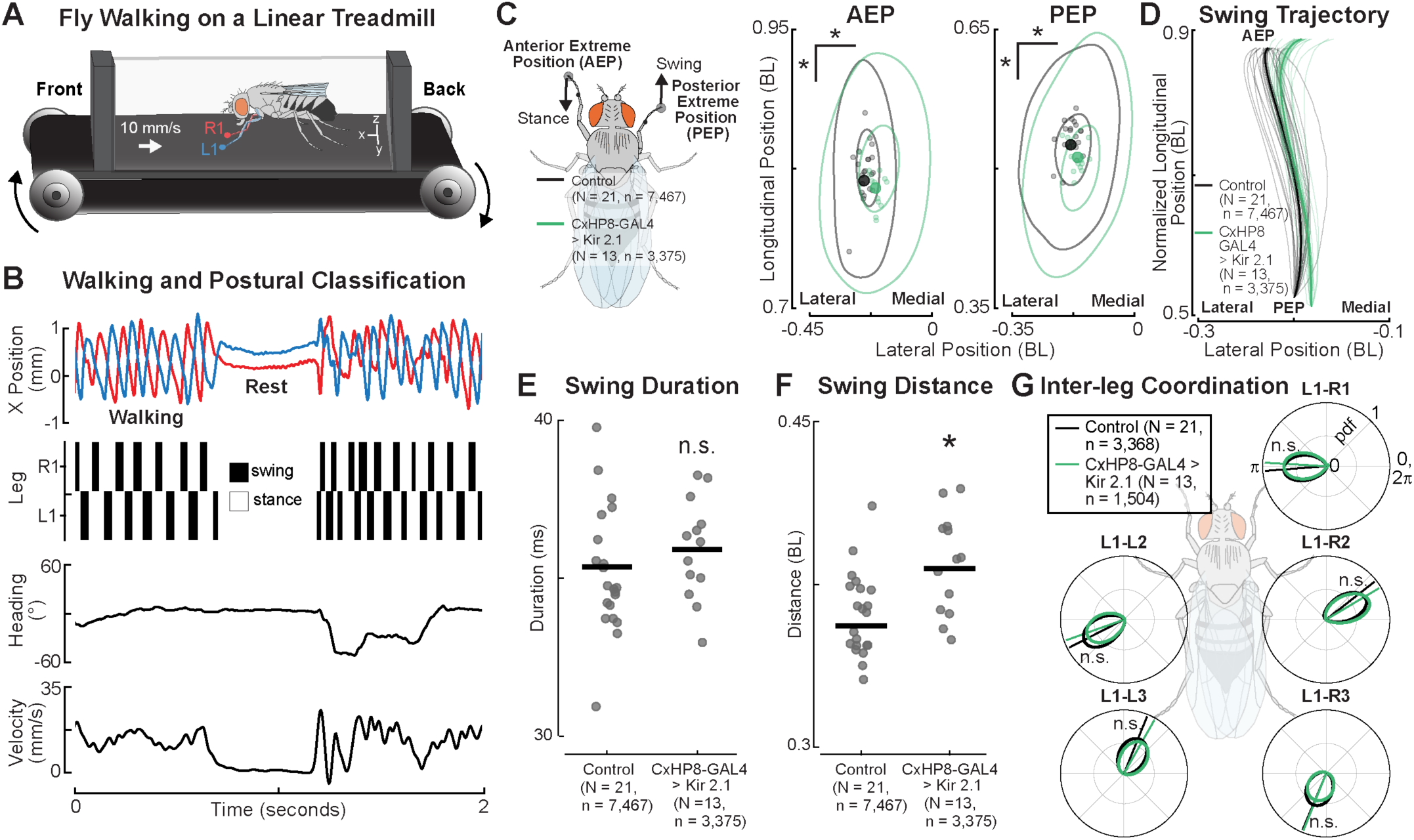
CxHP8 neurons control swing-to-stance transitions in unconstrained flies. (**A**) Schematic of the treadmill system. Flies were recorded with 5 high-speed cameras (200 fps) as they walked on a treadmill with a belt speed of 10 mm/s. The 3D positions of each distal leg tip and the head, thorax, and abdomen were tracked using automated pose estimation methods. (**B**) Example trial period illustrating that both walking kinematics and resting posture can be obtained using the treadmill system. Shown are the longitudinal (X) position and the classification of swing and stance of the left and right front leg as the fly walked and rested on the treadmill belt. Walking and rest periods were classified using the heading and body velocity of the fly. (**C**) Left: Schematic of the anterior and posterior extreme positions. Right: Kernel density estimation plots of the anterior and posterior extreme positions significantly shifted medially and posteriorly in flies with CxHP8 neurons chronically silenced (green) with the inward rectifying potassium channel, Kir 2.1. N: number of flies; n: number of steps. Bootstrapping with a Bonferroni correction of 12 was used to statistically compare CxHP8 silenced and control flies (*: p < 0.004). (**D**) Swing trajectories of CxHP8 silenced (green) and control (black) flies demonstrate the medial overshoot near the swing-to-stance transition when CxHP8 neurons were silenced. Each trajectory was scaled by the average swing distance and began at the average posterior extreme position. Solid line: mean trajectory; Thin lines: mean trajectory of each fly; Shaded region: 95% confidence interval; N: number of flies; n: number of steps. (**E**) Swing duration was not statistically different between CxHP8 silenced and control flies, based on a t-test with a Bonferroni correction of 2 on the means of flies (n.s.: p > 0.025). (**F**) Swing distance was significantly greater for CxHP8 silenced flies compared to controls (*: p < 0.025). (**G**) Inter-leg coordination (relative step cycle phase) between the left front leg and each other leg was not significantly different between CxHP8-silenced and control flies. A Kuiper two-sample test with a Bonferroni correction of 5 determined significance (n.s.: p > 0.01). N: number of flies; n: number of phase comparisons. See also **Figure S4**.

Like tethered flies, we also found silencing CxHP8 neurons in flies walking on the treadmill impaired step kinematics of the front legs near the swing-to-stance transition. In particular, the anterior extreme position (AEP), where a leg first enters stance, was displaced medially and posteriorly compared to controls (**Figure 4C**). The posterior extreme position (PEP), where a leg first enters swing, was also significantly displaced in a similar manner to the AEP. Given that the AEP marks the swing-to-stance transition, and that changes in the AEP may drive changes in the PEP, we focused on understanding the kinematic changes that facilitate the displacement of the AEP in CxHP8-silenced flies. In examining the average swing trajectories of the front legs, we found a significant medial deviation of the trajectory near the swing-to-stance transition in CxHP8 silenced flies compared to controls (**Figure 4D**). Although this deviation did not result from a significantly longer mean swing duration (**Figure 4E**), the mean swing distance was significantly greater in flies with CxHP8 neurons silenced (**Figure 4F**). Unlike front leg step kinematics, inter-leg coordination was not significantly altered in flies with CxHP8 neurons silenced (**Figure 4G**). Overall, the silencing of CxHP8 neurons in flies walking on the treadmill resulted in more medial front leg movement near the swing-to-stance transition, consistent with our results from tethered flies.

In larger insects, hair plates have been shown to be important for controlling resting body (Wendler, G., 1972) and leg posture (Bässler, 1977; Theunissen et al., 2014). We therefore examined the posture of flies during rest periods on the treadmill. Even though flies with CxHP8 neurons silenced didn’t have significantly different body heights (**Figure S4A**) or angles (**Figure S4B**), the resting positions of their tarsi were significantly more splayed out than controls (**Figure S4C**). We further quantified this by calculating the area of the polygon formed by the tarsi positions (**Figure S4D**). The mean polygon area of CxHP8 silenced flies was significantly greater than controls, providing further support that CxHP8 neurons control the resting posture of the legs.

### Connectome analysis enables functional predictions for other hair plates on the fly leg

Our calcium imaging data indicate that CxHP8 neurons function as limit detectors of anterior coxa movement (**Figure 1**). Based on analysis of the connectome, we hypothesized that feedback from CxHP8 neurons helps transition the coxa away from its anterior limit by driving posterior movement (**Figure 2**). We tested and confirmed this prediction by genetically manipulating the activity of CxHP8 neurons in behaving flies (**Figures 3,4**). Having validated the predicted reflex function for one hair plate, we now return to the connectome to predict the role of other front leg hair plates in leg motor control.

We focused on the two other hair plates on the front leg coxa, CxHP3 and CxHP4, along with the three hair plates located at the coxa-trochanter joint, TrHP5, TrHP6, and TrHP7 (**Figure 5A**). In FANC, we reconstructed the axons of each of these hair plates (**Figure 5B, S5A**). Of the thousands of output synapses from these hair plate axons (**Figure S5B**), the majority of synapses are onto local premotor and motor neurons (**Figure 5C**). Interestingly, we also found that some hair plate axons synapse onto glia, suggesting that their signaling may be used for non-motor related functions, such as detecting injury (**Figures 5C**, **S5C**). The axons of each hair plate connect to distinct populations of motor neurons that actuate the leg segment the hair plate is located on (**Figure 5D**). For instance, CxHP4 axons provide strong connectivity to motor neurons of the coxa rotate motor module, which drives anterior coxa movement, whereas the antagonistically positioned CxHP8, connects to the antagonistic coxa posterior motor module. We also found that the axons of each hair plate synapse onto diverse premotor neurons (**Figure S5D**). By combining the connectivity of hair plates onto premotor and motor neurons, we constructed 2-layer sensorimotor circuits for each hair plate (**Figure S5E**). We also computed the motor impact score for each hair plate (**Figure 5E**), a measure that weights direct and indirect connectivity between sensory and motor neurons, incorporating the neurotransmitter of any intervening interneurons (Lee et al., 2024). This impact score is useful to understand trends or differences in motor connectivity, but, importantly, does not consider factors such as activity dynamics or intrinsic neural properties. Nonetheless, comparing impact scores revealed clear differences in motor connectivity among hair plates on different parts of the leg. For example, our analysis predicts that CxHP4 drives anterior leg movement, CxHP3 reinforces both anterior and posterior leg movement, TrHP5 and TrHP7 drive posterior leg movement, and TrHP6 drives anterior leg movement. Overall, the connectivity of hair plates suggests each limit detector is specialized for controlling specific leg movements that are matched to the different extremes of joint movement that they detect.

**Figure 5.**
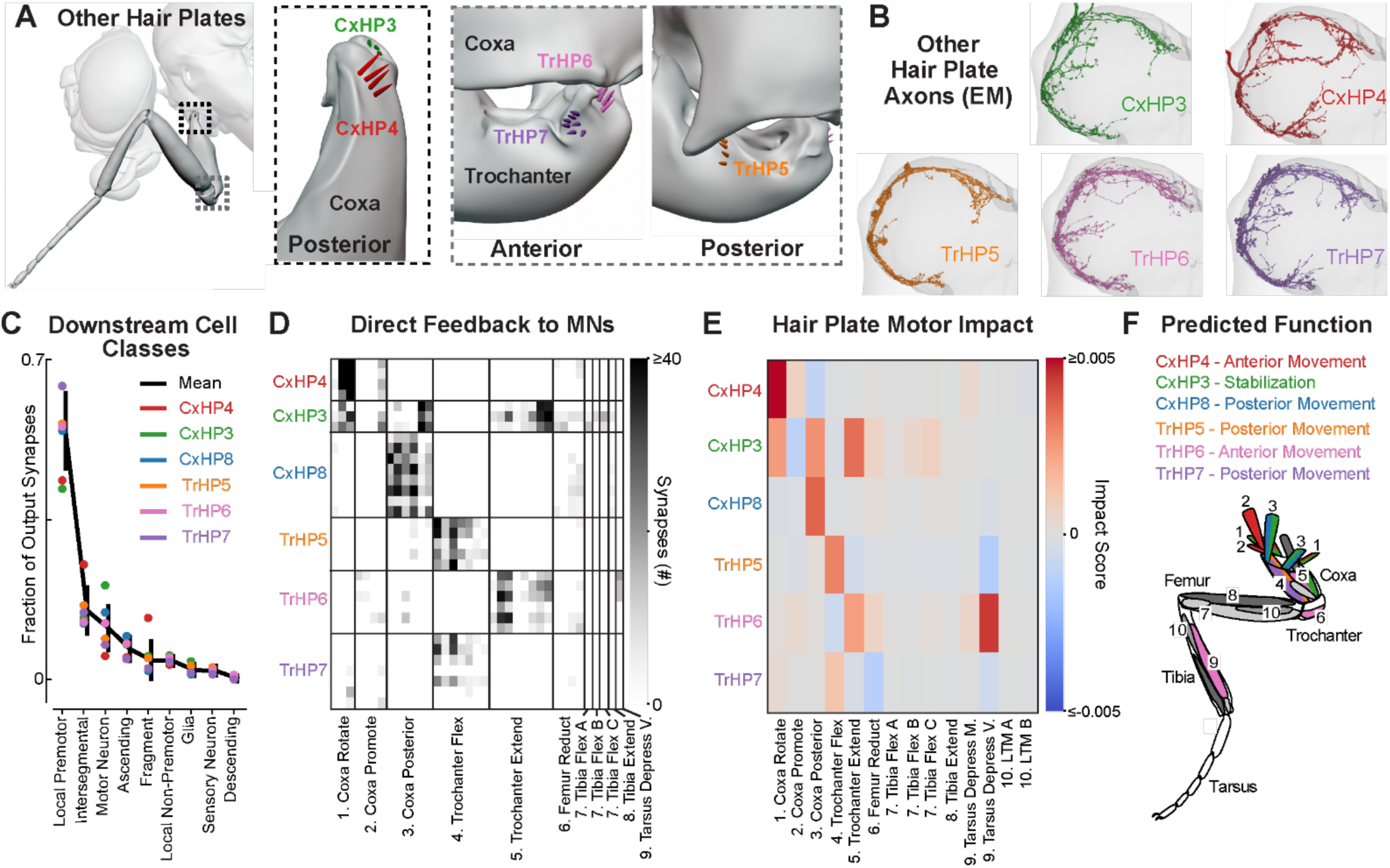
Synaptic connectivity of hair plate axons indicates distinct roles in leg motor control. (**A**) Hair plates located at the thorax-coxa (black dotted box) and coxa-trochanter (gray dotted box) joints of the front leg. Schematics were rendered in Blender and based on high-resolution confocal images. We focused on two hair-plates, in addition to CxHP8, located at the thorax-coxa joint (CxHP4: red; CxHP3: green) and three at the coxa-trochanter joint (TrHP5: orange; TrHP7: purple; TrHP6: pink). (**B**) Reconstructed hair plate axons within the left front leg neuromere of FANC. (**C**) Fraction of hair plate axon output synapses onto VNC cell classes. Black line: mean fraction of output synapses across all hair plate axons; Color dot: mean fraction of output synapses for the axons of each hair plate. (**D**) Connectivity matrix shows the number of synapses that motor neurons receive from hair plate axons. Axons from the same hair plate are grouped together (rows) while motor neurons within their defined motor module are arranged according to the proportion of input synapses they receive from specific hair plate axons. The leg muscles corresponding to the motor modules 1-9 are shown in **F**. (**E**) Motor impact scores (Lee et al., 2024), weighted sums of direct (monosynaptic) and indirect (di-synaptic) signed connection strengths, of hair plates onto leg motor modules. Red and blue hues correspond to excitatory and inhibitory impact scores, respectively. The leg muscles corresponding to the motor modules 1-10 are shown in **F**. See **Methods** for more information on how the motor impact score was calculated. (**F**) Predicted function and muscles controlled by each hair plate, as indicated by corresponding colors. See also **Figure S5** and **Video 2**.

## Discussion

In this study, we analyzed the physiology, circuit connectivity, and behavioral function of hair plates on the fly leg. We first used calcium imaging to discover that the neurons of one hair plate (CxHP8) are tonically active at the anterior extremes of coxa movement (**Figure 1**). We then used connectomics to predict that feedback from CxHP8 neurons drives posterior coxa and leg movement (**Figure 2**). Optogenetic stimulation of CxHP8 neurons was sufficient to drive reflexive posterior leg movements (**Figure 3**). We also found that optogenetic and chronic silencing of CxHP8 neurons in tethered flies (**Figure 3**) and flies walking freely on an actuated treadmill (**Figure 4**), caused the leg to overshoot the swing-to-stance transition in a medial direction. In addition to their role in walking, silencing CxHP8 neurons altered the resting position of the legs (**Figure S4**). Overall, these results agree with the sensorimotor reflex predictions from the connectome. We also provide functional predictions for the other hair plates on the front leg (**Figure 5**), which can be tested in future experiments as new genetic driver lines become available. Altogether, this work illustrates an integrated approach, anchored in the connectome, to infer the sensorimotor functions of spatially distributed proprioceptors.

Proprioceptive limit detectors are common in limbed animals. Nearly fifty years ago, recordings from joint receptors in the cat leg found that single afferents fired in a phasic or tonic manner as they neared the limit of the range of joint movement (Burgess and Clark, 1969). Previous electrophysiological work in locusts (BrÄunig and Hustert, 1985a, 1985b; Haskell, 1959; Kuenzi and Burrows, 1995; Newland et al., 1995) and cockroaches (French and Sanders, 1979; French and Wong, 1976; Pearson et al., 1976; Wong and Pearson, 1976) also found neurons within the same hair plate had either tonic or phasic encoding properties. However, we found no evidence for phasic tuning among CxHP8 axons – calcium signals remained sustained when the leg was held at an extreme position. It is possible that all hair plate neurons in the fly have tonic encoding properties. It is also possible that other hair plate neurons, which we did not record from in this study, respond phasically to mechanical stimulation. More work is needed to determine whether physiological properties vary across different hair plates.

While extensive prior work has investigated the encoding properties of proprioceptive limit detectors in other animals, less was known about their downstream connectivity and contributions to leg motor control during behavior. We found that activating and silencing CxHP8 neurons confirmed our leg motor control predictions based on connectivity. It is worth noting that the VNC circuits connecting hair plates and leg motor neurons are far more complex than our behavioral results and past work would suggest. For example, CxHP8 neurons also disynaptically excite and/or inhibit motor neurons controlling every other segment within the leg (**Figure 2B**), suggesting leg-wide rather than leg segment-specific motor control. Premotor neurons downstream of hair plates are also recurrently connected in complex patterns, and some synapse back onto CxHP8 axons (**Figure S2B**). This dense connectivity may support inter-joint coordination or provide robustness to external perturbations that the animal faces while navigating complex terrain.

Reflex circuits may also be modulated (i.e., suppressed or amplified) during different behavioral contexts. Hook (but not claw) proprioceptors in the fly femoral chordotonal organ (FeCO) are presynaptically inhibited during self-generated leg movements (Dallmann et al., 2023). Here, we did not observe suppression of hair plate calcium activity during spontaneous leg movements (**Figure 1**). Interestingly, optogenetic silencing of CxHP8 neurons had a significant, but relatively small, effect on extreme anterior extension of the coxa during front leg grooming (**Figure S3D**), even though CxHP8 neurons showed strong activity during this behavior (**Figures 1J**, **S1D, Video 3**). Gain modulation could occur in downstream premotor neurons to fine-tune the reflex action of hair plates.

The fruit fly’s front leg contains over 200 proprioceptive sensory neurons, including hair plates, campaniform sensilla, and chordotonal neurons (Kuan et al., 2020). Chordotonal neurons are primarily concentrated in the FeCO, which senses the femur/tibia joint (Mamiya et al., 2018). The more proximal coxa and trochanter joints contain similar numbers of campaniform sensilla and hair plates. Connectomic and electrophysiological evidence suggest that signals from these different proprioceptor classes are highly integrated in downstream circuits. For instance, work in the stick insect has shown that load and movement signals encoded by campaniform sensilla and FeCO neurons, respectively, are integrated in premotor networks to control the femur-tibia joint (Gebehart et al., 2021). Similarly, in the fly, electrophysiology and calcium imaging has revealed that signals from functional subtypes of FeCO neurons (i.e., claw, hook, club) are integrated in downstream neurons (Agrawal et al., 2020; Chen et al., 2021). Connectomic reconstruction has also revealed convergence of claw, hook, hair plate, and campaniform sensilla in downstream VNC neurons (Lee et al., 2024). Altogether, integration of signals across diverse proprioceptors may contribute to locomotor robustness and could also help explain why silencing CxHP8 neurons had only a very subtle effect on joint kinematics during walking.

Here, we focused on local reflex action of hair plates within a single leg. Hair plate axons do not project intersegmentally, and the majority of their disynaptic connectivity is also onto motor neurons controlling the same leg. However, past work has shown that hair plates may also contribute to intersegmental coordination across legs. In stick insects, for example, the ablation of hair plates on the middle leg resulted in non-linear and greater shifts in the anterior extreme position of the ipsilateral hind leg (Cruse et al., 1984). We did not observe kinematic changes of other legs when manipulating CxHP8 neurons on the fly’s left front leg. However, intersegmental effects could be driven by other hair plates.

Unlike other senses with a dedicated and localized organ like an eye, ear, or nose, proprioception is a distributed sense that relies on mechanosensory neurons distributed throughout the body. Therefore, a fundamental challenge for understanding proprioception is that the same class of sensory neurons in different locations may be anatomically specialized to detect specific perturbations to the body. Investigating hundreds or thousands of proprioceptors, one-by-one, is experimentally intractable. A promising future approach to overcome this problem is to combine the methods used above (connectomics, calcium imaging, and genetic manipulations during behavior) with biomechanical modeling (Vaxenburg et al., 2024; Wang-Chen et al., 2023). To facilitate this effort, we have added all of the hair plates characterized in this study to the open-source Janelia fly body Blender model (Vaxenburg et al., 2024) (**Video 2**). As other components are added to the body model, it will become possible to simulate hair plate signaling through recurrent, connectome-constrained networks. In the future, integrating biomechanical and connectome-constrained network models may provide a promising solution to the challenge of understanding how distributed sensory systems, including proprioception, control complex behaviors.

## Acknowledgements

We thank members of the Tuthill and Brunton Labs for technical assistance and feedback on the manuscript. We thank Seok Jun Moon for sharing the CxHP8 split-GAL4 driver line (R48A07 AD: R20C06 DBD). B.G.P was supported by an NSF Graduate Research Fellowship (Fellow ID: 2018261272). C.J.D was supported from the Deutsche Forschungsgemeinschaft (DFG, German Research Foundation) project 432196121. Other support was provided by National Institutes of Health grants R01NS102333, R01NS128785, and U19NS104655, a Searle Scholar Award, a Klingenstein-Simons Fellowship, a Pew Biomedical Scholar Award, a McKnight Scholar Award, a Sloan Research Fellowship, the New York Stem Cell Foundation, and a UW Innovation Award to J.C.T. J.C.T is a New York Stem Cell Foundation – Robertson Investigator.

## Author Contributions

B.G.P. and J.C.T. conceived the study and wrote the manuscript. I.S. created the gorgeous Blender model of a fruit fly with hair plates, as well as Video 2. B.G.P and A.S. performed confocal imaging of hair plates. B.G.P. and A.C. proofread the hair plate axons and their downstream and upstream partners in FANC. C.J.D. collected and processed the calcium imaging data for CxHP8. G.M.C. and S.W.B. collected optogenetic activation and silencing datasets for CxHP8. B.G.P. analyzed the connectivity, calcium imaging, and optogenetic datasets. B.G.P collected and analyzed the kinematics and posture of flies walking in the treadmill setup. A.A. provided valuable feedback about the connectivity of hair plates onto motor circuits.

## Lead contact

Further information and requests for resources and reagents should be directed to and will be fulfilled by the lead contact, John C. Tuthill.

## Data and Code Availability

Data is available on Dryad (DOI: 10.5061/dryad.fxpnvx153). Code for analyzing and visualizing hair plate connectivity in the EM dataset, calcium activity of CxHP8 neurons, kinematics during optogenetic experiments, and walking kinematics and posture during treadmill experiments is located on GitHub (https://github.com/Prattbuw/Hair_Plate_Paper).

## Supplemental information

**Figure S1.**
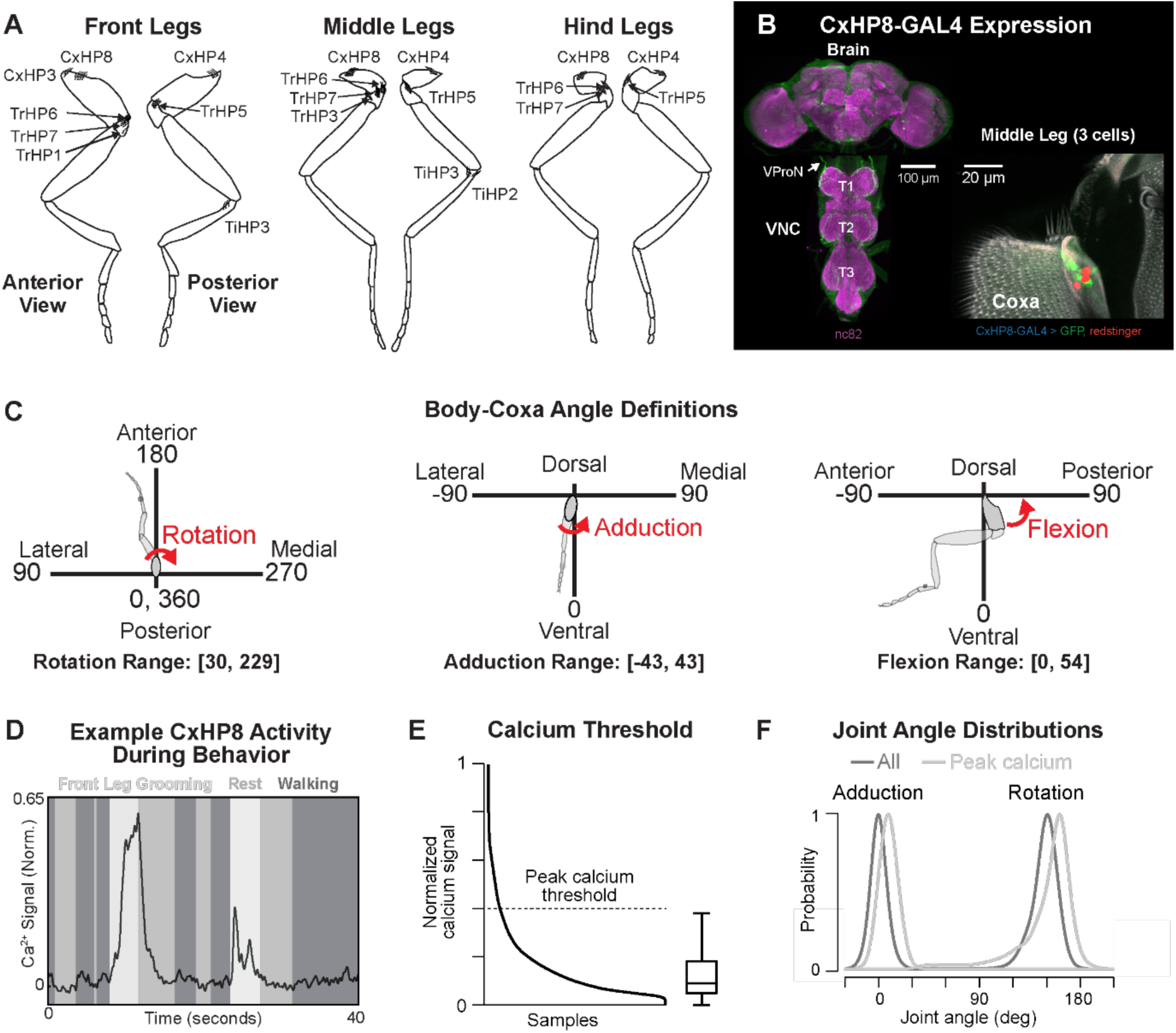
Hair plate locations on the legs of *Drosophila*, CxHP8 driver line expression, and thorax-coxa joint angle definitions and encoding. (**A**) Schematics of the front, middle, and hind legs showing the locations of hair plates. (**B**) Confocal images showing the expression of the CxHP8 split-GAL4 in the middle leg (3 of 8 neurons labeled), brain, and VNC. The ultrastructure (autofluorescence: gray) of CxHP8 consists of 8 cuticular hairs located on the coxa. Labeled mechanosensory neuron (GFP: green) project through the ventral prothoracic nerve (VProN) into the VNC (nc82: magenta). The cell bodies of CxHP8 neurons are labeled by the nuclear marker, redstinger (red). (**C**) Reference frame for the rotation, adduction, and flexion thorax-coxa angles of the front leg. The measured range of each angle is displayed at the bottom. (**D**) Example trial showing the normalized calcium activity (normalized to the max calcium activity in the dataset) during front leg grooming, rest, and walking. (**E**) Calcium threshold was set at the upper whisker limit (Q3 + 1.5 * IQR) of the calcium activity box plot and was used for determining large peak calcium activity events. (**F**) Adduction and rotation joint angle probability distributions during all calcium activity (gray) and just peak calcium events (light gray).

**Figure S2.**
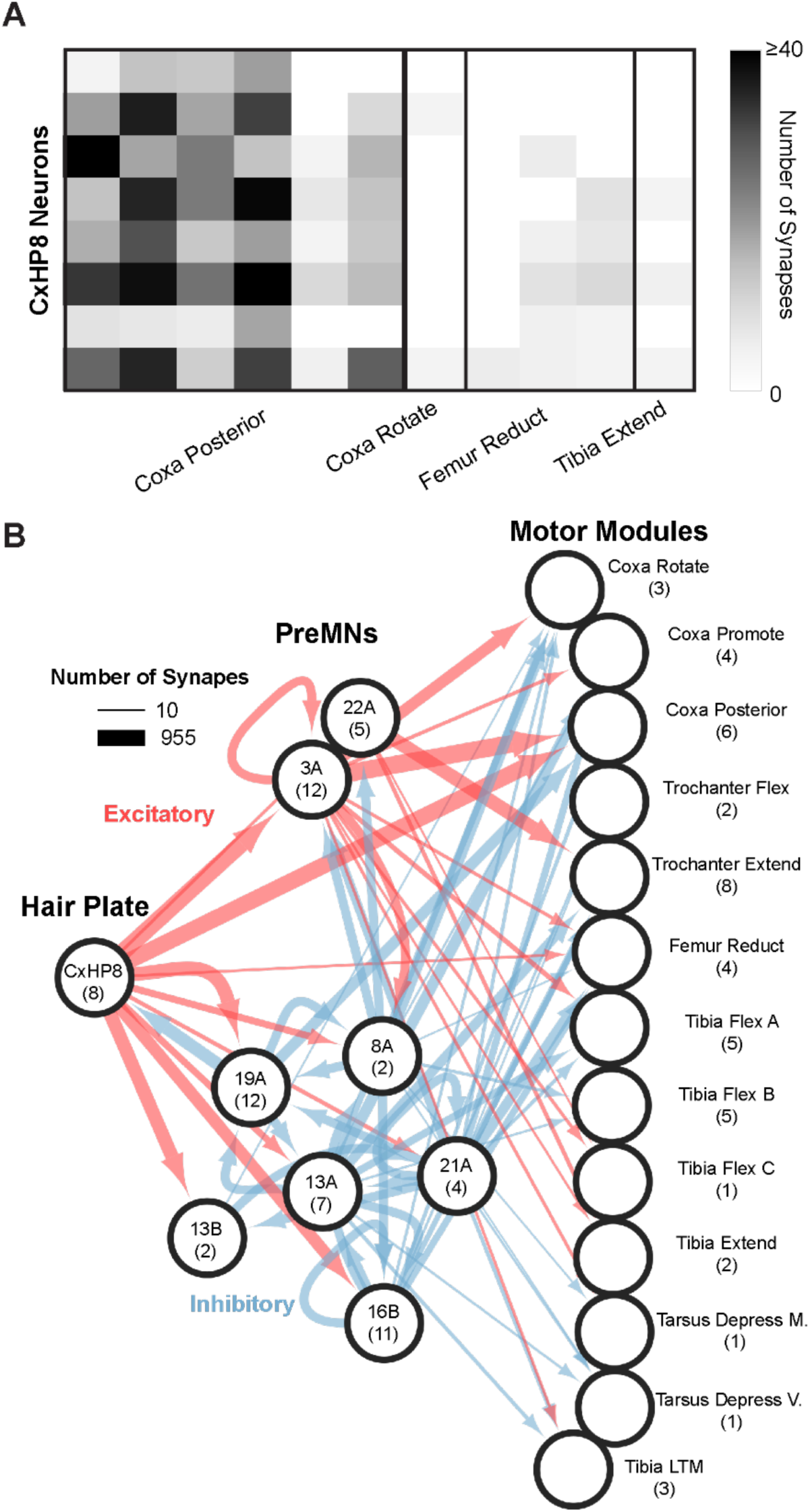
CxHP8 reflex circuit connectivity. (**A**) CxHP8 connectivity onto motor neurons reveals that all CxHP8 axons primarily synapse onto motor neurons within the coxa posterior motor module. The x-axis represents individual motor neurons within defined motor modules. (**B**) CxHP8 reflex circuit shown in Figure 2B, with the addition of premotor-to-premotor, and premotor-to-CxHP8 recurrent connections. Red and blue lines indicate excitatory and inhibitory connections, respectively. The width of the line corresponds to the number of synapses between cell classes. The number of neurons within each class are indicated within the parentheses. Note that only connections with greater than 3 synapses on average between upstream neurons within a class and downstream cell classes were used in this analysis.

**Figure S3.**
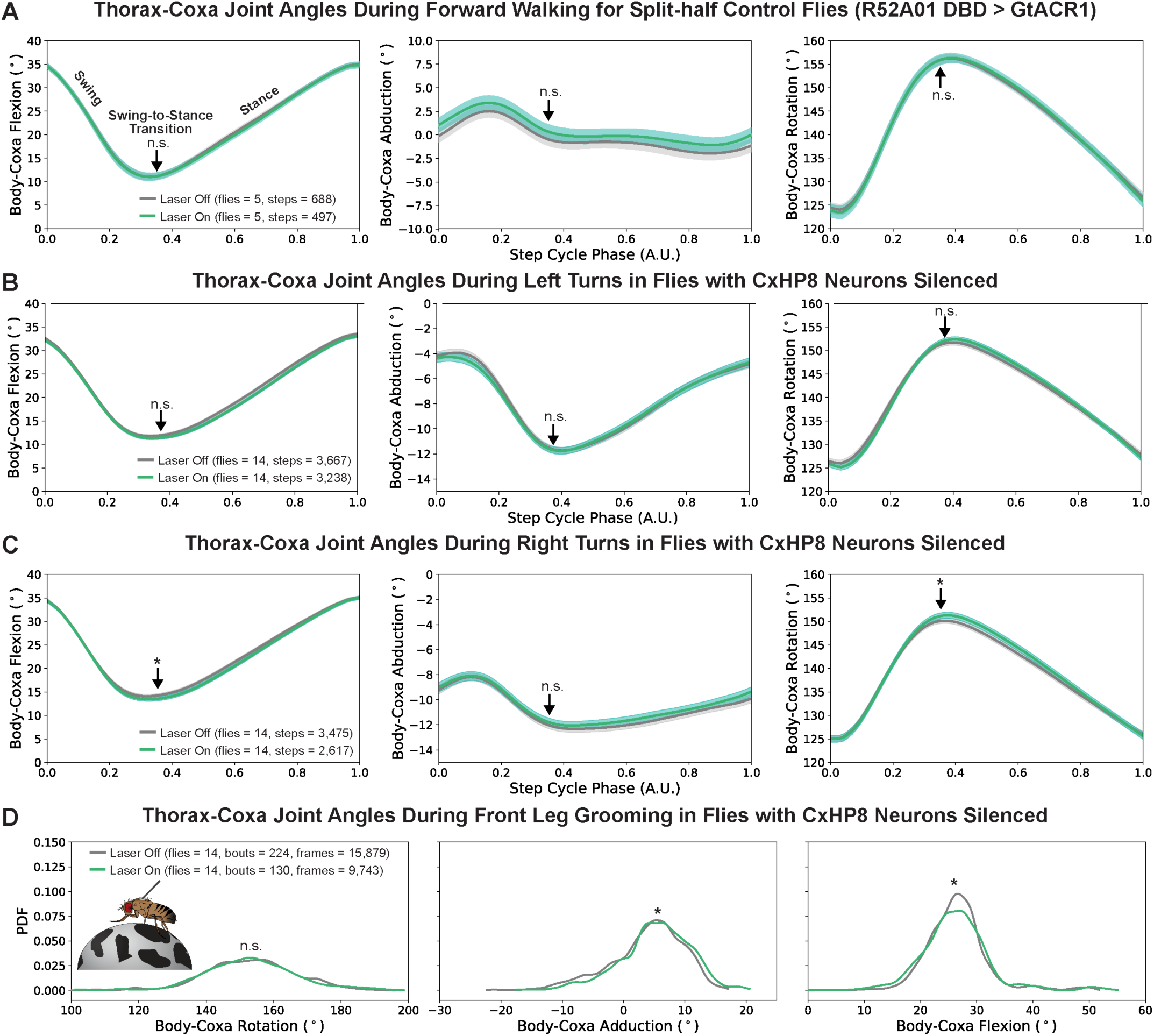
Control flies had no significant change in joint angles during laser stimulation and CxHP8 silencing had significant effects on thorax-coxa joint angles during right turns and front leg grooming but not left turns. (**A**) Control flies (R52A01 DBD > GtACR1) had no significant (n.s.: p > 0.025) changes in coxa joint angles when the green laser was on compared to when it was off. Arrow: the average swing-to-stance transition of the step cycle; Shaded region: 95% confidence interval; N: number of flies; n: number of steps. A t-test with a Bonferroni correction of 2 was used to statistically compare the distributions of joint angles at the swing-to-stance transition. (**B**) Optogenetic silencing (green) of CxHP8 neurons in flies turning left didn’t have a significant effect (t-test with a Bonferroni correction of 2; n.s.: p > 0.025) on thorax-coxa angles compared to turning steps when the laser was off. (**C**) Optogenetic silencing of CxHP8 neurons had a significant (t-test; *: p < 0.025) effect on flexion and rotation angles at the swing-to-stance transition during right turns. (**D**) Optogenetic silencing (green) of CxHP8 neurons significantly (t-test with a Bonferroni correction of 2; *: p < 0.025) altered the thorax-coxa adduction and flexion joint angle during front leg grooming compared to when the laser was off (gray).

**Figure S4.**
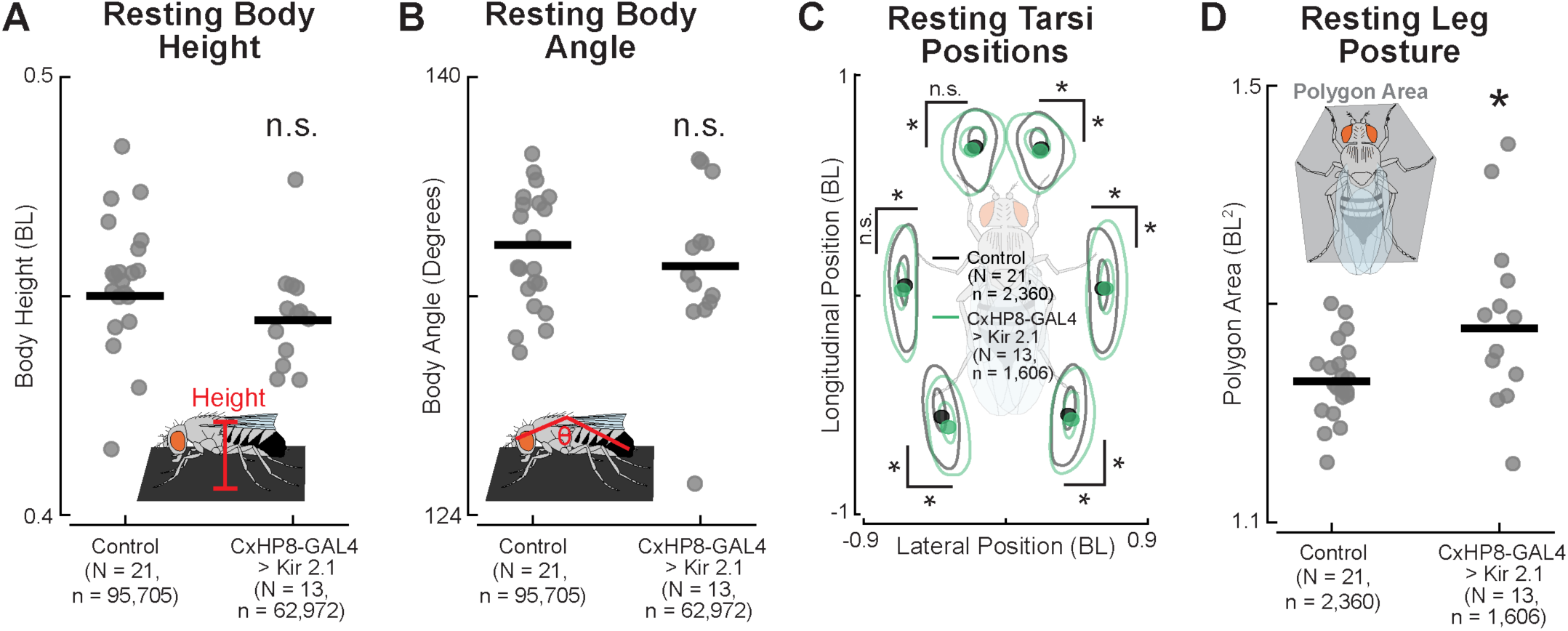
CxHP8 feedback controls resting leg positions but not body height or angle. (**A**) Body height was lower on average but not significantly different between CxHP8 silenced and control flies (t-test; p > 0.05). N: number of flies; n: number of frames. (**B**) Body angle was also smaller on average but not significantly different between CxHP8 silenced and control flies (t-test; p > 0.05). N: number of flies; n: number of frames. (**C**) Resting positions of the tarsi during periods of standing were significantly different between CxHP8 silenced (green) and control (black) flies. Distributions were determined by kernel density estimation. Dot: mean position; N: number of flies; n: number of resting bouts. Bootstrapping with a Bonferroni correction was used to test for statistical differences (*: p < 0.001). (**D**) As quantified by polygon area, the resting posture of the legs was significantly more spread out for flies with CxHP8 neurons silenced compared to controls (t-test; *: p < 0.05). N: number of flies; n: number of resting bouts.

**Figure S5.**
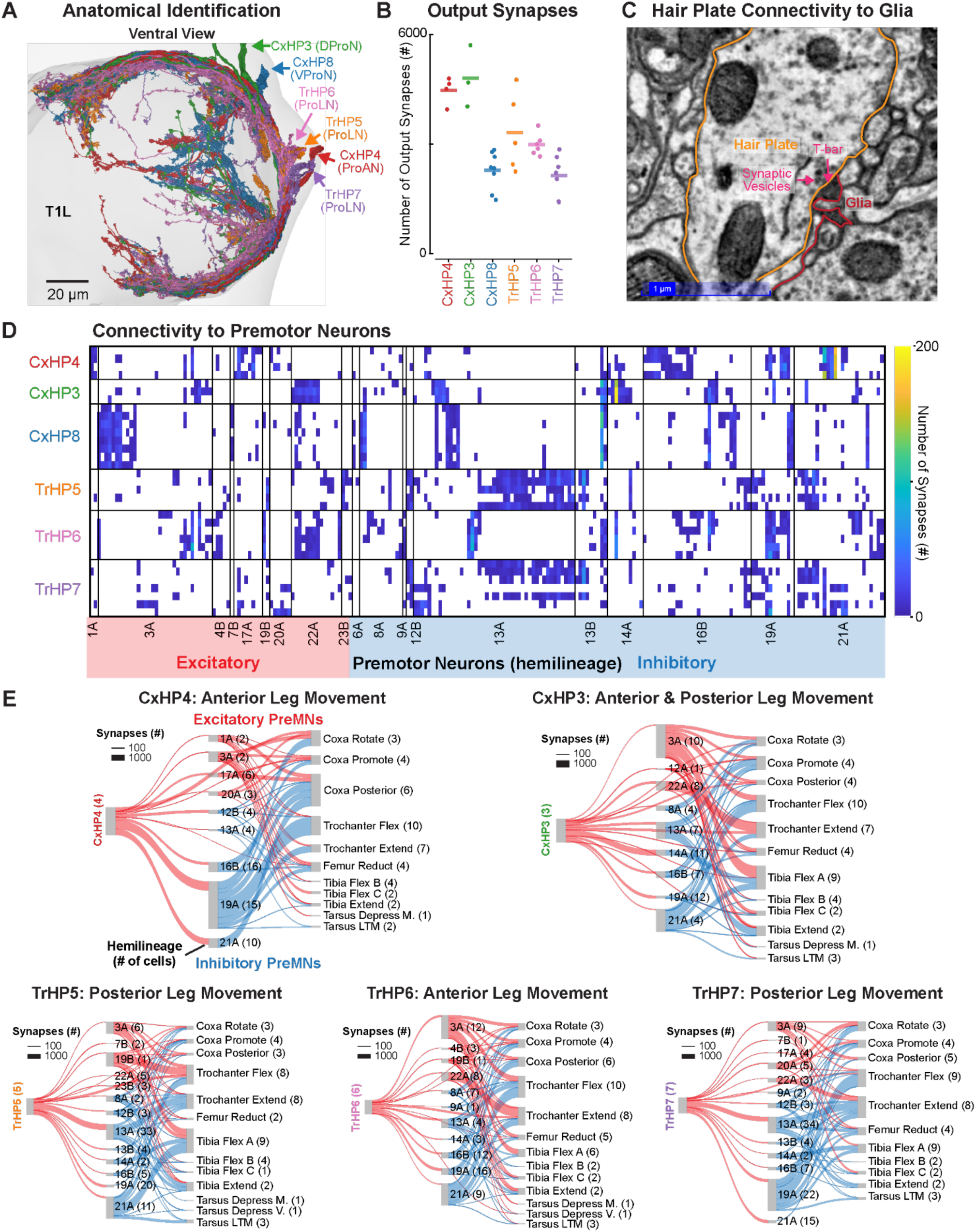
Synaptic connectivity of hair plates in motor circuits and onto glia. (**A**) Front leg hair plate axons are identified, in part, based on the nerve they project through and axonal morphology. DProN: dorsal prothoracic nerve; VProN: ventral prothoracic nerve; ProAN: prothoracic accessory nerve; ProLN: prothoracic leg nerve. (**B**) Total number of predicted output synapses of hair plate axons in FANC. Line: mean output synapses; dot: number of output synapses of a single hair plate axon. (**C**) Electron microscopy image illustrating the connectivity between a hair plate axon (orange) and a glia cell (red). Synaptic vesicles and a putative T-bar are labeled. (**D**) Connectivity matrix showing the number of synapses that hair plate axons supply to premotor neurons. Premotor neurons are classified as excitatory (red) or inhibitory (blue) based on their identified hemilineage. Premotor neurons within a hemilineage are arranged according to the proportion of input synapses they receive from each hair plate axon. (**E**) Based on their connectivity with premotor and motor neurons, the neurons of each hair plate are predicted to engage reflex circuits that control distinct aspects of leg movement. Above each reflex circuit is the prediction for the associated hair plate in controlling leg movement. Only connections of at least 4 synapses on average per neuron within a cell class onto downstream cell classes were used to construct each reflex circuit. The number of cells in each cell class is indicated within the parentheses.

**Table S1.** Proprioceptor expression of split-GAL4 driver lines in the legs of *Drosophila melanogaster*. The number of cells labeled by the screened split-GAL4 driver lines was determined by counting cells genetically expressing membrane bound GFP and a nuclear reporter (i.e. UAS-mcD8GFP; Redstinger) using confocal imaging. Proprioceptors that did not have their expression checked received a “N/A”. The lack of a column for a particular proprioceptive class indicates that either no expression was seen or that proprioceptor was not checked for expression. HP: hair plate; CS: campaniform sensilla.

| Driver Line | Front Leg Proprioceptors |  |  |  |  |  |  |  |  |  | Middle Leg Proprioceptors |  |  |  |  |  |  | Hind Leg Proprioceptors |  |  |  |  |  |
| --- | --- | --- | --- | --- | --- | --- | --- | --- | --- | --- | --- | --- | --- | --- | --- | --- | --- | --- | --- | --- | --- | --- | --- |
|  | CxHP4 | CxHP3 | CxHP8 | TrHP5 | TrHP6 | TrHP7 | TrHP1 | TrCS3 (TrG) | TrCS5 (TrFa) | FeCS11 | CxHP4 | CxHP8 | TrHP5 | TrHP6 | TrHP7 | TrCS3 | TrCS2 | CxHP4 | CxHP8 | TrHP5 | TrHP6 | TrHP7 | TrCS3 |
| R48A07 AD:<br>R20C06 DBD | 0 | 0 | 6 | 0 | 0 | 0 | 0 | 0 | 0 | 0 | 0 | 3 | 0 | 0 | 0 | 0 | 0 | 0 | 0 | 0 | 0 | 0 | 0 |
| R93D09 AD:<br>VT061711 DBD | 2 | 0 | 4 | 3 | 0 | 0 | 0 | 3 | 0 | 0 | 0 | 6 | 3 | 3 | 1 | 2 | 0 | 4 | 6 | 3 | 4 | 2 | 0 |
| R93D09 AD:<br>VT009832 DBD | 2 | 2 | 0 | 3 | 2 | 2 | 0 | 0 | 0 | 0 | 2 | 0 | 3 | 2 | 0 | 0 | 1 | 3 | 0 | 3 | 3 | 0 | 0 |
| R93D09 AD:<br>VT040055 DBD | 1 | 0 | 0 | 2 | 0 | 0 | 0 | 2 | 0 | 0 | 2 | 0 | 0 | 0 | 0 | 2 | 0 | 2 | 0 | 0 | 0 | 0 | 2 |
| R93D09 AD:<br>R37G07 DBD | 0 | 0 | 0 | 2 | 1 | 1 | 0 | 0 | 0 | 0 | 1 | 0 | 1 | 0 | 0 | 0 | 0 | 2 | 0 | 0 | 3 | 2 | 0 |
| VT061711 AD:<br>VT045603 DBD | 4 | 0 | 5 | 2 | 0 | 0 | 0 | 3 | 0 | 0 | 3 | 6 | 3 | 2 | 0 | 2 | 0 | 3 | 8 | 2 | 4 | 3 | 3 |
| R93D09 AD:<br>VT038115 DBD | 0 | 0 | 0 | 2 | 2 | 0 | 0 | 0 | 0 | 0 | 0 | 0 | 2 | 0 | 0 | 0 | 0 | 0 | 0 | 2 | 0 | 0 | 0 |
| VT038115 AD:<br>VT040055 DBD | 0 | 0 | 0 | 2 | 0 | 3 | 0 | 0 | 0 | 0 | N/A | N/A | N/A | N/A | N/A | N/A | N/A | N/A | N/A | N/A | N/A | N/A | N/A |
| VT0098323 AD:<br>VT061711 DBD | 1 | 0 | 0 | 2 | 0 | 2 | 0 | 2 | 0 | 0 | N/A | N/A | N/A | N/A | N/A | N/A | N/A | N/A | N/A | N/A | N/A | N/A | N/A |
| VT0098323 AD:<br>R20C06 DBD | 0 | 0 | 0 | 0 | 0 | 2 | 0 | 3 | 0 | 1 | N/A | N/A | N/A | N/A | N/A | N/A | N/A | N/A | N/A | N/A | N/A | N/A | N/A |
| R39B11 AD:<br>VT061711 DBD | 0 | 0 | 0 | 3 | 0 | 2 | 0 | 2 | 0 | 3 | N/A | N/A | N/A | N/A | N/A | N/A | N/A | N/A | N/A | N/A | N/A | N/A | N/A |
| R48A07 AD:<br>VT040055 DBD | 0 | 0 | 0 | 1 | 0 | 1 | 1 | 0 | 0 | 0 | N/A | N/A | N/A | N/A | N/A | N/A | N/A | N/A | N/A | N/A | N/A | N/A | N/A |
| VT038115 AD:<br>VT061711 DBD | 0 | 0 | 0 | 2 | 0 | 2 | 0 | 0 | 0 | 0 | N/A | N/A | N/A | N/A | N/A | N/A | N/A | N/A | N/A | N/A | N/A | N/A | N/A |
| VT0098323 AD:<br>R94E06 DBD | 1 | 0 | 0 | 0 | 0 | 0 | 0 | 0 | 0 | 1 | N/A | N/A | N/A | N/A | N/A | N/A | N/A | N/A | N/A | N/A | N/A | N/A | N/A |
| R22D12 AD:<br>VT061711 DBD | 2 | 0 | 0 | 0 | 1 | 1 | 0 | 3 | 2 | 3 | N/A | N/A | N/A | N/A | N/A | N/A | N/A | N/A | N/A | N/A | N/A | N/A | N/A |
| R22D12 AD:<br>R94E06 DBD | 2 | 0 | 4 | 0 | 1 | 0 | 0 | 0 | 0 | 1 | N/A | N/A | N/A | N/A | N/A | N/A | N/A | N/A | N/A | N/A | N/A | N/A | N/A |
| VT038115 AD:<br>R94E06 DBD | 0 | 0 | 0 | 0 | 0 | 0 | 0 | 0 | 0 | 0 | N/A | N/A | N/A | N/A | N/A | N/A | N/A | N/A | N/A | N/A | N/A | N/A | N/A |
| R48A07 AD:<br>R22D12 DBD | 4 | 0 | 0 | 0 | 3 | 0 | 0 | 0 | 0 | 0 | N/A | N/A | N/A | N/A | N/A | N/A | N/A | N/A | N/A | N/A | N/A | N/A | N/A |
| R94E06 AD:<br>VT040055 DBD | 1 | 0 | 0 | 0 | 0 | 3 | 0 | 0 | 0 | 0 | N/A | N/A | N/A | N/A | N/A | N/A | N/A | N/A | N/A | N/A | N/A | N/A | N/A |

## Video Legends

**Video 1. Calcium activity of CxHP8 neurons while the left front leg was moved passively and slowly using a 3-axis platform.**

**Video 2. Animation of hair plates on the front leg of a fly walking on a treadmill.** The fly and hair plates were rendered in Blender and based on high resolution confocal images. Depicted are the three hair-plates located at the thorax-coxa joint (CxHP4: red; CxHP3: green; CHP8: blue) and three at the coxa-trochanter joint (TrHP5: orange; TrHP7: purple; TrHP6: pink).

**Video 3. Calcium activity of CxHP8 neurons while a fly behaved on a spherical treadmill.**

## Methods

### Fly Husbandry and Genotypes

Experiments were performed on adult male flies, *Drosophila melanogaster*, between 2-5 days post-eclosion. Flies were reared in a 25°C incubator with 14:10 light:dark cycle within vials filled with a standard cornmeal and molasses medium. Genotypes used in experiments are listed in Table 1.

**Table 1.** *Drosophila melanogaster* genotypes used for experiments.

| Stock Name | Genotype | Figure(s) | Stock Source |
| --- | --- | --- | --- |
| R48A07 AD: R20C06 DBD > mCD8GFP; Redstinger | w[1118]/yw, hs flp, UAS-mCD8-GFP; P{+t[7.7]w[=mC]=R48A07-p65.AD}attP40/ UAS-stingerRed; P{y[+t7.7]w[+mC]=R20C06-GAL4.DBD}attP2/+ | 1, S1 | UAS-mCD8GFP; Redstinger was gifted from the Parrish lab, University of Washington and crossed with split-GAL4 gifted from Moon Lab, Yonsei University (Bloomington Stocks #69841 and #71070) |
| R48A07 AD: R20C06 DBD > pJFRC7-20XUAS-IVS-mCD8::GFP | w[1118]/w[*]; P{+t[7.7]w[=mC]=R48A07-p65.AD}attP40/ +; P{y[+t7.7]w[+mC]=R20C06-GAL4.DBD}attP2/P{pJFRC7-020XUAS-IVS-mCD8::GFP}attP2 | 1, S1 | 20X UAS-mCD8::GFP was gifted from the Rubin lab and crossed with split-GAL4 gifted from Moon Lab, Yonsei University (Bloomington Stocks #69841 and #71070) |
| R48A07 AD: R20C06 DBD > GcaMP7f; tdTomato | w[1118]/+ DL; P{+t[7.7]w[=mC]=R48A07-p65.AD}attP40/ P{w[+mC] = UAS-tdTom.S}2; P{y[+t7.7]w[+mC]=R20C06-GAL4.DBD}attP2/ PBac{y[+mDint2]w[+mC]=20xUAS-IVS-jGCaMP7f}VK00005 | 1, S1 | UAS-GCaMP7f; tdTomato were contrusted from Bloomington stocks #36327 and #79031, and crossed with split-GAL4 gifted from Moon Lab, Yonsei University (Bloomington Stocks #69841 and #71070) |
| R48A07 AD: R20C06 DBD > ChromsonR (CxHP8 optogenetic activation) | w[1118]/+ DL; P{+t[7.7]w[=mC]=R48A07-p65.AD}attP40/+ DL; P{y[+t7.7]w[+mC]=R20C06-GAL4.DBD}attP2/10X UAS ChromsonR mCherry (attp2)/TM3 Sb | 3, S3 | UAS-ChrimsonR was gifted from Janelia (outcrossing was done by Anne Sustar, University of Washington) and crossed with split-GAL4 gifted from Moon Lab, Yonsei University (Bloomington Stocks #69841 and #71070) |
| R48A07 AD: R20C06 DBD > GtACR1 (CxHP8 optogenetic silencing) | w[1118]/+ DL; P{+t[7.7]w[=mC]=R48A07-p65.AD}attP40/+ DL; P{y[+t7.7]w[+mC]=R20C06-GAL4.DBD}attP2/20X-UAS-GtACR1-EYFP (III) attp2 | 3, S3 | UAS-GtACR1 stock was created by Anne Sustar in the Tuthill Lab at the University of Washington and was crossed with split-GAL4 gifted from Moon Lab, Yonsei University (Bloomington Stocks #69841 and #71070) |
| R52A01 DBD > GtACR1 (light-effect control) | w[1118]/+ DL; +/+ DL; P{y[+t7.7]w[+mC]=R52A01-GAL4.DBD}attP2/20X-UAS-GtACR1-EYFP (III) attp2 | S3 | UAS-GtACR1 stock was created by Anne Sustar in the Tuthill Lab at the University of Washington and was crossed with Bloomington Stock #69141 |
| R48A07 AD: R20C06 DBD > Kir 2.1 (CxHP8 chronic silencing) | w[1118]/+ DL; P{+t[7.7]w[=mC]=R48A07-p65.AD}attP40/+ DL; P{y[+t7.7]w[+mC]=R20C06-GAL4.DBD}attP2/pJFRC49-10XUAS-IVS-eGFP::Kir2.1(attP2) | 4, S4 | Crossed Dickinson Lab Stock U-111 with split-GAL4 gifted from Moon Lab, Yonsei University (Bloomington Stocks #69841 and #71070) |
| R48A07 AD > Kir 2.1 (Control) | w[1118]/+ DL; P{+t[7.7]w[=mC]=R48A07-p65.AD}attP40/+ DL; pJFRC49-10XUAS-IVS-eGFP::Kir2.1(attP2)/+ | 4, S4 | Crossed Dickinson Lab Stock U-111 with Bloomington Stock #71070 |
| R39B11 AD > Kir 2.1 (Control) | w[1118]/+ DL; P{+t[7.7]w[=mC]=R39B11-p65.AD}attP40/+ DL; pJFRC49-10XUAS-IVS-eGFP::Kir2.1(attP2)/+ | 4, S4 | Crossed Dickinson Lab Stock U-111 with Bloomington Stock #71040 |

### Blender Model of *Drosophila* Body and Front Leg Hair Plates

A morphologically accurate model of an adult *Drosophila melanogaster* containing the hair plates at the thorax-coxa and coxa-trochanter joints of the front leg was constructed using the 3D rendering and modeling software, Blender (Vaxenburg et al., 2024). The body of the fly, and the anatomy and location of hair plates in the model were based on high-resolution confocal images. We replayed the 3D walking kinematics of the left front leg coxa measured from a tethered fly walking on the ball to this model to illustrate the phases during the step cycle when the hairs of each coxa hair plate would be deflected (**Video 2**).

### Confocal Imaging of Proprioceptor Expression

#### Peripheral expression of proprioceptors in legs

Front, middle, and/or hind legs of male flies associated with the driver lines in **Table S1** crossed with the UAS-mcD8GFP; redstinger reporter line were fixed in a 4% formaldehyde (PFA) PBS solution for 20 minutes and then washed three times in a 0.2% Triton X-100 PBS solution. Legs were then cleared with FocusClear (CelExplorer) and mounted on slides in MountClear (CelExplorer). The expression at each hair plate was assessed using an Olympus FV1000 confocal microscope. The number of proprioceptor cells labeled by each driver line was then determined through visual inspection of the volumetric imaging stacks in FIJI (Schindelin et al., 2012)

#### Central expression of proprioceptors in the brain and VNC

The brain and VNC were dissected from male flies associated with the split-GAL4 driver lines that showed interesting peripheral expression in hair plates, such as the CxHP8 split-GAL4 driver line (i.e. R48A07 AD; R20C06 DBD). Unlike above, the GAL4 driver resulted in the expression of pJFRC7-20XUAS-IVS-mCD8::GFP. The brains and VNCs were fixed in the same manner as legs but then were put into a blocking solution (5% goat serum, PBS, 0.2% Triton-X) for 20 minutes, followed by being incubated in a blocking and primary antibody solution (1:50 concentration of anti-GFP chicken antibody, 1:50 concentration of anti-brp mouse) for 24 hours at room temperature. The brains and VNCs were then washed three times in PBS with 0.2% Triton-X and incubated in a blocking and secondary antibody solution (1:250 concentration of anti-chicken-Alexa 488, 1:250 concentration of anti-mouse-Alexa 633). After this, the brains and VNCs were washed three times with PBST and mounted on a slide in Vectashield (Vector Laboratories). The brains and VNCs were imaged with the same Olympus confocal microscope as legs and image stacks were analyzed in FIJI (Schindelin et al., 2012).

### Reconstruction and Identification of Hair Plate Axons in FANC EM dataset

#### Hair Plate Axon Reconstruction and Connectivity

Hair plate axons were reconstructed in the electron microscopy dataset of the female ventral nerve cord, FANC (Azevedo et al., 2024; Phelps et al., 2021), through manual proofreading of the automatically segmented cell fragments in Neuroglancer (Maitin-Shepard et al., 2021). Annotations containing information about the reconstructed hair plate axons, such as what hair plate they project from, we uploaded to the Connectome Annotation Versioning Engine (CAVE) and are publicly available to those that have access to the FANC dataset. We then determined and proofread the cell fragments that received or provided at least 3 synapses from or onto hair plate axons. In addition to proofreading the upstream and downstream partners of hair plates, we classified them, if possible, into general cell types and specific cell classes using pre-existing CAVE annotation tables and/or expert knowledge (Azevedo et al., 2024; Harris et al., 2015; Lacin et al., 2019; Lesser et al., 2024). The annotations of these neurons were also uploaded to CAVE and are publicly available. We determined the downstream and upstream partners of hair plates and performed connectivity analyses using a custom Python script. Note that only neurons that had 4 or more output synapses on average per neuron within a cell class onto individual downstream neurons were used in the reflex circuit analyses.

#### Hair Plate Identification

Hair axons were identified from previously traced backbones and annotations in CATMAID (Kuan et al., 2020; Phelps et al., 2021). The axons of CxHP8, CxHP3, and CxHP4 were identified based on if they projected through the ventral prothoracic, dorsal prothoracic, or prothoracic accessory nerves, respectively (**Figure S5A**). All trochanter hair plate axons project through the prothoracic leg nerve (**Figure S5A**), so TrHP5, TrHP6, and TrHP7 neurons were instead identified based on their location in the nerve bundle, axonal morphology, and downstream connectivity with motor neurons (**Figure 5D**). TrHP6 axons were identified because they make distinct connections onto motor modules compared to TrHP5 and TrHP7 (**Figure 5D**). TrHP5 and TrHP7 axons could be differentiated because they enter the leg neuropil through the ProLN at different locations (**Figure S5A**).

#### Motor Impact Score

Motor impact score (Lee et al., 2024) was computed by first calculating the monosynaptic weight between a hair plate and motor module, which is the total number of synapses that all neurons within a hair plate make onto the motor neurons within a motor module, divided by the total number of synapses onto that motor module across all neurons. Next, we calculated the di-synaptic weight between a hair plate and motor module by finding and summing the proportion of hair plate input synapses provided to each premotor neuron connected to the motor module of interest, multiplied by the proportion of input synapses those premotor neurons make onto the motor module. Note that the fractional input between these premotor neurons and the motor module is signed, based on the hemilineages of the premotor neurons. The motor impact score is then derived by summing the monosynaptic and di-synaptic weights.

### Calcium Imaging of Hair Plate Axons

We used a previously described preparation, setup, and analysis pipeline (Dallmann et al., 2023) to image calcium signals in hair plate axons in the neuromere of the left front leg during active and passive leg movements.

#### Preparation

We attached a fly to a custom-made holder, removed a rectangular piece of cuticle from the dorsal thorax, removed the underlying indirect flight muscles, and used an insect pin to displace the digestive system. This provided a dorsal view on the hair plate axons entering the VNC from the left front leg.

#### Two-photon image acquisition

Calcium signals in hair plate axons were recorded with a two-photon Movable Objective Microscope (MOM; Sutter Instruments) with a 20x water-immersion objective (Olympus XLUMPlanFI, 0.95 NA, 2.0 mm wd; Olympus). Hair plate neurons expressed the calcium indicator GCaMP7f (green fluorescence) and the structural marker tdTomato (red fluorescence). Fluorophores were excited at 920 nm by a mode-locked Ti:sapphire laser (Chameleon Vision S; Coherent). We used a Pockels cell to keep the power at the back aperture of the objective below ~35 mW. Emitted fluorescence was directed to two high-sensitivity GaAsP photomultiplier tubes (Hamamatsu Photonics) through a 705 nm edge dichroic beamsplitter followed by a 580 nm edge image-splitting dichroic beamsplitter (Semrock). Fluorescence was band-passed filtered by either a 525/50 (green) or 641/75 (red) emission filter (Semrock). Image acquisition was controlled with ScanImage 5.2 (Vidrio Technologies) in Matlab (MathWorks). The microscope was equipped with a galvo-resonant scanner, and the objective was mounted onto a piezo actuator (Physik Instrumente; digital piezo controller E-709). We acquired volumes of three 512 x 512 pixel images spaced 5 μm apart in depth (10 μm total) at a speed of 8.26 volumes per second.

#### Two-photon image analysis

Two-photon images were smoothed with a Gaussian filter (sigma = 3 pixels; size = 5 x 5 pixels). Each tdTomato image was aligned to the average tdTomato signal of the recorded trial using a cross-correlation-based image registration algorithm (Guizar-Sicairos et al., 2008). The same alignment was used for the GCaMP images. We averaged the three GCaMP and tdTomato images per volume. Then, we extracted the mean fluorescence in manually drawn regions of interest (ROIs). To correct for vertical movement of the VNC, we computed the ratio of GCaMP fluorescence to tdTomato fluorescence in each frame. To facilitate comparisons across trials and flies, ratio values were z-scored by subtracting the mean of a baseline ratio and dividing by the standard deviation of that baseline ratio. The baseline was defined in each trial as the 10% smallest ratio values. Finally, z-scored ratio values were upsampled to the sampling rate of leg tracking (300 Hz) using cubic spline interpolation and then low-pass filtered using a moving average filter with a time window of 0.2 s.

#### Platform and treadmill

The platform consisted of a metal pin (0.5 mm diameter, 4.4 mm length) mounted onto a three-axis micromanipulator (MP-225; Sutter Instruments). The pin was wrapped in black sandpaper to provide sufficient grip for the flies’ tarsi. The micromanipulator was controlled manually.

The treadmill consisted of a patterned Styrofoam ball (9.1 mm diameter; 0.12 g) floating on air in an aluminum holder. The air flow was set to ~500 ml/min. The ball was illuminated by two infrared LEDs (850-nm peak wavelength; ThorLabs) via optical fibers. Ball movements were recorded at 30 Hz with a camera (Basler acA1300-200um; Basler AG) equipped with a macro zoom lens (Computar MLM3X-MP; Edmund Optics). Ball rotations around the fly’s cardinal body axes (forward, rotational, sideward) were reconstructed offline using FicTrac(Moore et al., 2014). Rotational velocities of the fly were calculated based on the known diameter of the ball. Velocities were upsampled to the sampling rate of leg tracking (300 Hz) using cubic spline interpolation and low-pass filtered using a moving average filter with a time window of 0.2 s.

#### Leg tracking

Movement of the left front leg was recorded at 300 Hz with two cameras (Basler acA800-510um; Basler AG) equipped with 1.0x InfiniStix lenses (68 mm wd; Infinity) and 875 nm short pass filters (Edmund Optics). The leg was illuminated by an infrared LED (850-nm peak wavelength; ThorLabs) via an optical fiber. We used a previously trained (Dallmann et al., 2023) deep neural network (Mathis et al., 2018) to automatically track all leg joints in each camera view. 2D tracking data from both camera views were then combined to reconstruct leg joint positions and angles in 3D using Anipose (Karashchuk et al., 2021).

#### Behavior classification

Fly behavior was classified semi-automatically based on thresholds on the velocity of the front and middle leg tarsi as described previously (Dallmann et al., 2023). Flies on the platform were classified as resting or actively moving. Flies on the treadmill were classified as resting, walking, or grooming. All classifications were reviewed and manually corrected if necessary.

#### Data selection

Frames were manually excluded from the analysis if the front leg was involved in movements other than walking or grooming on the treadmill (e.g., extended downward pushing), the femur-tibia joint of the front leg was not tracked correctly, or the two-photon image registration failed (e.g., the VNC moved out of the imaging volume).

### Optogenetic Activation and Silencing Experiments

Optogenetic experiments were performed on adult male flies that were raised on 35mM in 95% EtOH ATR for 1-3 days, were 2-5 days old, de-winged, and fixed to a rigid tether (0.1 mm thin tungsten rod) with UV glue (KOA 300). These flies were placed onto a spherical foam ball (weight: 0.13 g; diameter: 9.08mm) suspended by air within a visual arena. A dark bar with a width of 30 degrees with respect to the fly oscillated at 2.7 Hz in front of the fly. A spatially precise red (638 nm; 1200 Hz pulse rate; 30% duty cycle, Laserland) or green (532 nm; 1200 Hz pulse rate; 60% duty cycle, Laserland) laser was then positioned on the thorax-coxa joint of the left front leg. Optogenetic activation experiments were conducted on flies in which their CxHP8 neurons expressed ChrimsonR, whereas silencing experiments were performed on those that expressed GtACR1 in those neurons (**Table 1**). Trials were 2 seconds in duration and consisted of turning the laser on (experimental trials) or keeping it off (control trials) 0.5 seconds into the trial for 1 second. During each trial, the behavior each fly was recorded with 6 high-speed cameras (300 fps; Basler acA800-510 µm; Balser AG) and the movement of the ball was recorded at 30 fps with a camera (FMVU-03MTM-CS) and processed using FicTrac (Moore et al., 2014). The 3D positions of each leg joint and the corresponding joint angles were determined by using DeepLabCut (Mathis et al., 2018) and Anipose (Karashchuk et al., 2021). Kinematic analyses were performed in a custom Python script. Behaviors were classified using a previously published random forest classifier (Karashchuk et al., 2021). In addition, front leg grooming was also classified based on the proximity and velocity of the front legs.

### Treadmill Experiments

Treadmill experiments were performed on 2-5 day old CxHP8 silenced and genetically-matched control male flies (**Table 1**) using a previously described treadmill system and experimental protocol (Pratt et al., 2024). The inward rectifying potassium channel, Kir 2.1, was used to silence the activity of CxHP8 neurons throughout development. In short, trials consisted of placing a de-winged fly into a chamber and moving the belts of the split-belt treadmill at a steady-state speed of 10 mm/s for 20 minutes in total. During which, we recorded the behavior of the fly at 200 fps and in 20 second bouts with 5 high-speed cameras (Basler acA800-510 µm; Balser AG). We then used DeepLabCut (Mathis et al., 2018) and Anipose (Karashchuk et al., 2021) to reconstruct the 3D positions of head, thorax, abdomen, and each leg tip. Custom Python scripts were used to analyze and visualize walking kinematics and posture.

### Kinematic Classification and Parameters

The classification and quantification of behavior, swing and stance, and most walking kinematic parameters for tethered flies and flies walking on the treadmill were conducted in the same manner as a prior study (Pratt et al., 2024). For tethered flies in both the calcium imaging and optogenetic experiments, steps or frames were classified as forward walking if the instantaneous forward velocity and absolute rotational velocity of the ball was greater than 5 mm/s and less than 25 degrees/s, respectively. Left and right turns were classified as periods where the rotational velocity was less than −25 degrees/s or greater than 25 degrees/s, respectively. Other behaviors, like rest/sanding and front leg grooming, were determined by using a previously published behavioral classifier (Karashchuk et al., 2021). For flies walking on the treadmill, forward walking was classified as periods where the body velocity had a forward velocity greater than 5 mm/s and the absolute heading angle with respect to the front of the chamber was less than 15 degrees. Rest periods are classified as those where the instantaneous velocity of the body and all tarsi was less than 5 mm/s (i.e. all legs in stance).

During walking, the classification of each leg into swing and stance was based on the smoothed (i.e. with a Gaussian Kernel) instantaneous velocity of the tarsus of each leg. Instantaneous values were negative during periods where the tarsus moved posteriorly along the longitudinal axis of the body. A leg was classified as being in swing if its tarsus had an instantaneous velocity greater than 5 mm/s or less than −25 mm/s. Otherwise, the leg was classified as being in stance. Short, 1 video frame, classifications of swing or stance were corrected and reclassified as the other phase of the step cycle.

Classified forward walking steps were further filtered to remove steps that occurred during the transition into forward walking or produced kinematics that were inconsistent with what was previously published (DeAngelis et al., 2019; Mendes et al., 2013; Strauss and Heisenberg, 1990; Wosnitza et al., 2013). Any step that didn’t meet the following criteria for a forward walking step was removed: step frequency between 5 and 20 steps/s, a swing duration between 15 and 75 ms, and a stance duration less than 200 ms. Steps that passed these filtering criteria were used for further kinematic analyses and comparisons.

All kinematic analyses and visualizations were done in custom Python scripts, which a publicly available on GitHub (https://github.com/Prattbuw/Hair_Plate_Paper).

### Statistical Analyses

Bootstrapping or t-tests were performed to determine statistical significance. If multiple comparisons were performed, a Bonferroni correction was employed to adjust the threshold of significance. t-tests were used to compare posture and step kinematics between CxHP8 silenced and control flies, along with joint angle distributions and tarsus positions during behavior. A Kuiper two-sample was used to assess statistical significance for inter-leg coordination. Finally, bootstrapping was used to statistically compare the anterior and posterior extreme positions between these flies.

### Glossary of Kinematic Parameters

**Body length (BL)**: the distance between the head and distal part of the abdomen.

**Body height**: the vertical distance between the ground and thorax.

**Body angle**: the angle made by the body frame.

**Anterior extreme position**: the position where a leg first contacts the ground (i.e. stance onset) in egocentric coordinates.

**Posterior extreme position**: the position where a leg first takes off from the ground (i.e. swing onset) in egocentric coordinates.

**Swing distance**: the total lateral and longitudinal distance a leg travels during the swing phase of the step cycle.

**Swing duration**: the duration of the aerial phase of leg movement during walking.

**L1 relative phase**: the relative offset in the stance onsets between the left front leg and the leg of interest with respect to the left front leg’s step cycle.

**Polygon area**: The area in BL^2^ of the polygon formed by the tarsi tip positions of the legs in stance.

